# Myofiber reconstruction at micron scale reveals longitudinal bands in heart ventricular walls

**DOI:** 10.1101/2022.05.12.491149

**Authors:** Drisya Dileep, Tabish A. Syed, Tyler F. W. Sloan, Perundurai S. Dhandapany, Kaleem Siddiqi, Minhajuddin Sirajuddin

## Abstract

The coordinated contraction of myocytes drives the heart to beat and circulate blood. Due to the limited spatial resolution of whole heart imaging and the piecemeal nature of high-magnification studies, a confirmed model of myofiber geometry does not yet exist. Using microscopy and computer vision we report the first three-dimensional reconstruction of myofibers across entire mouse ventricular walls at the micron scale, representing a gain of three orders of magnitude in spatial resolution over the existing models. Our analysis reveals prominent longitudinal bands of fibers that are orthogonal to the well-known circumferential ones. Our discovery impacts present understanding of heart wall mechanics and electrical function, with fundamental implications for the study of diseases related to myofiber disorganization.

## Introduction

The mammalian heart wall is densely packed with cardiomyocytes aligned end-on-end to form myofibers. The geometric organization of these fibers is critical for synchronous contraction, mechanical durability and electrical conduction (*1–4*). Malformations of myofibers can lead to pathological conditions of the heart, including cardiomyopathies, myocardial infarction and disorders related to electrical propagation (*5–7*). At a coarse spatial scale, the myofibers have been described as a helical continuum, wrapping around the chambers of the heart (*1, 2*). Several competing models of myofiber organization exist (*2, 8*), providing a conflicting picture. Thus, heart wall fiber geometry remains a fundamental unaddressed question in organ biology. Many of the models have been derived from millimeter resolution diffusion-tensor magnetic resonance imaging (DT-MRI) (*2, 9*– *12*). Such accounts suffer from limitations in spatial resolution since hundreds of cardiomyocytes can occupy a single voxel at this scale. Recovering myofiber geometry at the scale of individual cardiomyocytes is only possible using micron-scale light microscopy. However, studies in this direction have focused on small sections (*13*) or 3D stacks of heart tissue (*14, 15*) and therefore have neither recovered myofiber nor cardiomyocyte orientation at the whole heart scale. On the other hand, high resolution approaches based on histology are typically limited to two-dimensional imaging with the spatial positioning of the tissue samples being left unrecorded (*8, 16*). Inferring whole heart myofiber geometry from such histological recordings is thus not possible (*2*).

The geometric organization of myofibers across the heart wall is critical for its mechanical and electrical function, but it has not been understood at the scale of individual cardiomyocytes. To solve this fundamental problem, we combined confocal light microscopy based deep and wide imaging with computer vision techniques. This combination enabled us to obtain micron-scale cardiomyocyte orientations across entire long-axis and short-axis ventricular sections of mouse hearts. Our three-dimensional reconstructions at unprecedented spatial resolution revealed, for the first time, longitudinal bands of fibers extending along the entire length of the ventricular walls, orthogonal to the circumferential myofibers. Our findings represent a conceptual breakthrough in the understanding of heart myofiber geometry, thus opening new opportunities to investigate the structure of heart wall tissue and diseases affecting its function.

## Results

### Reconstruction of myocyte and myofiber orientation at the micron scale

The intact cleared hearts were serially sectioned, stained with fluorescent wheat germ agglutinin (WGA) and subjected to confocal imaging to cover an entire section using individual fields of view (Methods) (Figure. 1 and Supplementary Figure. 1). The images were further processed to recover micron scale resolution for each section and then used to estimate orientation at the individual myocyte scale using a structure-tensor method (*17*) (Methods) (Figure. 1 and Supplementary Figure. 2). This allowed us to recover cardiomyocyte orientation (represented as glyphs) at a higher resolution than previous studies, as well as myofiber organization (represented as streamlines) at a coarser spatial scale, spanning the entire section of heart tissue (Methods) (Figure. 1). In order to better analyze the estimated cell orientation fields, we plotted them using three angles: Theta (θ), Phi (Φ), and the Helix angle (αΗ) (Methods) (Figure. 2a). θ represents the acute angle between the projection of the estimated cell orientation (myofiber orientation) onto the short-axis (XY) plane and the X-axis. Φ represents the acute angle between the estimated cell orientation and the short-axis (XY) plane in the heart tissue. αΗ is the angle between the projection of the myofiber onto the local tangent plane to the heart wall and the Z-axis direction (Figure. 2a).

**Figure. 1:**
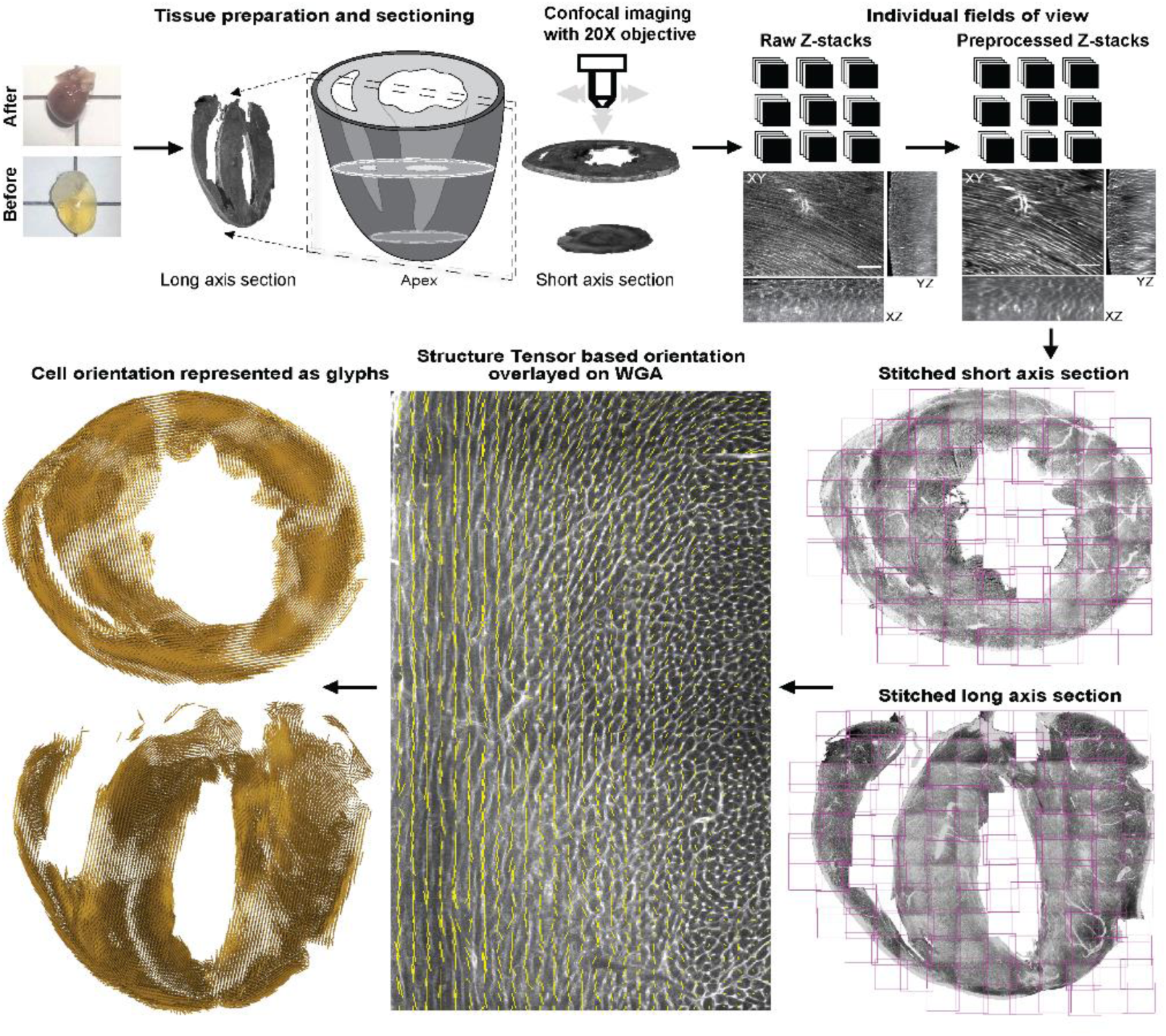
Overview of the heart tissue preparation, imaging and analysis pipeline. *Top panels from left to right;* Representative photographs of a mouse heart before and after applying the CLARITY procedure (*29*). An illustration of the cleared mouse heart sections analyzed in this study. The long-axis (LA) sections correspond to longitudinal dissections of the mouse heart through a transverse plane equivalent to the HLA-4C views of the AHA (or echocardiogram) nomenclature, revealing the right and left ventricular chambers. The short-axis (SA) sections are dissected at the mid-ventricular plane, equivalent to PSAX-PML views of the AHA (or echocardiogram) nomenclature (*34*). At least one mouse heart in this study provided four continuous sections, each being approximately 300 microns in thickness, for both the LA and SA analyses (Supplementary Table 1) (Methods). After sectioning and WGA staining, the tissue slices were imaged using confocal microcopy spanning the entire length, breadth and width of the LA and SA sections (Methods). Using custom built algorithms, the individual confocal stacks were preprocessed for contrast enhancement, denoised and then stitched (Methods). *Bottom panels from right to left;* The imaging and preprocessing pipeline results at a 2-micron voxel isometric resolution of the entire SA and LA sections, up to a depth of about 300 microns in thickness. A representative image of the WGA stain overlayed with the cell orientations estimated using a structure tensor method (Methods). The structure tensor eigenvectors associated with the smallest eigenvalue (yellow dashed lines) align with the long-axis of the cardiomyocytes. The estimated structure tensor vectors are visualized as glyphs represented as bold yellow lines for the entire SA and LA sections (Methods).

**Figure. 2:**
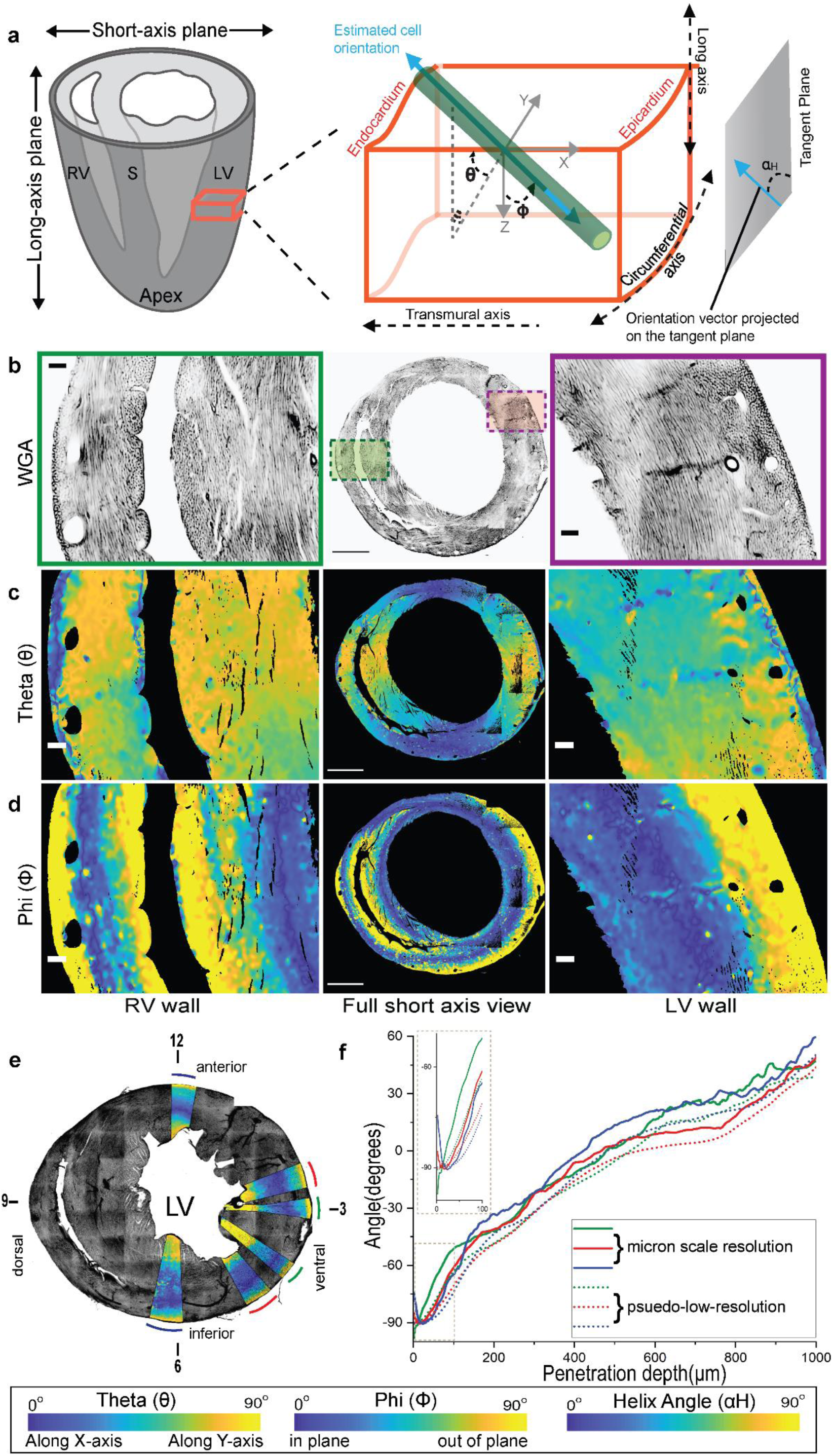
Cell orientation across a mid-ventricular short-axis section of the heart. **a.** An illustration of the angles measured to represent the cell orientations across the ventricular walls. The red box represents a magnified view of the ventricle wall, with the global axes and labels as indicated. The green cylinder and the blue bi-directional arrow represent the cardiomyocyte and its long-axis, i.e., the estimated cell orientation, respectively. Φ is the angle between the global Z-axis and the estimated cell orientation. θ is the angle between the projection of the estimated cell orientation onto the XY plane and the X-axis direction. αΗ, the helix angle, is the angle between the projection of the cell orientation onto the plane perpendicular to the transmural penetration direction and the circumferential direction. **b-d.** A magnified view of the right (green rectangle) and the left (magenta rectangle) ventricular regions for a full view of the WGA stain (middle). The θ and Φ angles for the mouse SA sections are shown using a parula colormap. The yellow tones for the θ angle represent cell orientation along the global Y-Axis, while the blue tones represent cell orientation along the X-axis. For Φ, the yellow and blue tones represent cells in-plane and orthogonal to the global Z-axis, respectively. The colormaps scale with the angles, as indicated. The scale bar for B is 1000 microns and for C and D is 100 microns. **e.** The individual 10° pie shaped sectors were combined into three groups; anterior/inferior, mid-ventral and adjacent to mid-ventral, representing the 12 O’clock and 6 O’clock, 3 O’clock and 4 O’clock, and 2 O’clock and 5 O’clock regions, respectively. The pie-sectorαΗ values were plotted and are highlighted as αΗ colormaps on a maximum intensity Z-projection of the WGA-stained short-axis section shown in grayscale. **f.** The average αΗ values* from micron-scale and pseudo-resolution analyses of the selected pie-shaped sectors of the LV region from one representative data set of a short-axis section (SAS3) is shown to the right. The αΗ values are plotted as a function of the penetration depth, depicted as distance in microns along the X-axis from the outer to the inner walls. * The values plotted here derive from combining two sectors, spanning 10 degrees in each sector selected, as illustrated.

### Regimes of discrete cell orientations across ventricular walls

A striking feature of the WGA-stained raw images is the appearance of discrete cell arrangements across the ventricular walls (Figure. 2a). To demarcate the boundaries between these different cell arrangements, we visualized the Φ and θ angles defined in Figure. 2a, using quantitative colormaps (Methods). The colormaps for the θ angle show that the component of the myofibers in the short-axis (XY) plane is perpendicular to the radial direction and follows a smooth continuum (Figure. 2c). Our imaging and analysis at the micron scale is consistent with previous reports that the myofibers wind around the ventricles in a circumferential manner (*2, 11, 20, 24, 25, 26*).

In contrast to θ, the Φ colormaps reveal a discrete and significant change in cardiomyocyte orientation across the ventricular walls (Figure. 2d). The out-of-plane cardiomyocytes, shown in yellowish tones, form a crescent shape along the outer walls of both ventricles and an orbicular shape in the inner walls surrounding both the chambers (Figure. 2d and Supplementary Figures 5, 6 and 7). Taken together, these reconstructions show that these out-of-plane cardiomyocytes are aligned to form continuous bands of longitudinal fibers that are orthogonal to the well-established circumferential myofibers.

### Sharp changes in myocyte orientation at ventricular wall boundaries

In a short-axis section, the Φ and αΗ angles are related to myofiber orientation with respect to the viewing plane. αΗ has been widely used to capture orientation changes along a transmural penetration from the outer to the inner heart wall (*2, 20, 21, 23*). We therefore computed αΗ at the micron scale using a ventricular outer boundary-based estimate of the radial penetration direction at each location in the left ventricular wall (Methods) (Supplementary Figure. 4, Figure. 2a). The average αΗ values for several pie-shaped sectors are plotted as a function of penetration depth (Supplementary Figures 9 and 10).

The patterns of αΗ values appear to be very similar at fixed angular distances away from the 3 O’clock position (Figure. 2e and Supplementary Figure. 9 a-d). Sectors in the vicinity of the 3 O’clock position on the short-axis section reveal a sharp drop of about 20 degrees within the first 50 microns near the circumference of the outer wall. Immediately after this sharp decline, αΗ undergoes a gradual increase of about 180 degrees for the remainder of the left ventricular wall, in a manner that is linearly proportional to penetration depth (Figure. 2e and Supplementary Figures 9 and 10). This latter smooth transition of αΗ through the myocardium (middle wall) until the endocardium (inner wall) is consistent with earlier findings (*2, 12, 18, 21–23*). However, the initial sharp transition in αΗ, closer to the epicardium (outer wall) is an entirely new discovery (Figure. 3B, C and Supplementary Figures 9 and 10). We hypothesize that earlier studies lacked the required spatial resolution to identify this sharp transition in αΗ. To test this, we averaged the structure tensor, estimated at the micron scale, to a coarser spatial resolution, resulting in a pseudo-low-resolution estimate of myofiber orientation (Methods). This orientation estimate is qualitatively equivalent to the millimeter or sub-millimeter scale of DT-MRI studies. A comparison of our micron scale and pseudo low-resolution αΗ angle estimates indeed shows the disappearance of the sharp change near the outermost layer, at the simulated coarser resolution (Figure. 2e). The longitudinal myofiber layer we have discovered in the outer ventricular wall might have remained obscure in past studies (*2, 12, 18, 21–23*) due to its narrow width of only approximately 50 microns. As such, in our simulated sub-millimeter scale coarse voxel analysis, it essentially disappears in the orientation plots (Figure. 2e). Our reconstructions also demonstrate the presence of narrow longitudinal myofiber layers in the inner ventricular chamber and septum walls (Figure. 2e).

### Charting the continuity of the longitudinal myofibers

The analysis of short-axis sections allowed us to identify narrow bands of myofibers near the boundaries of the ventricular walls that are aligned to the long-axis direction of the heart. To determine the spatial extent of these myofibers across the entire length of the ventricular walls, we turned our attention to the analysis of long-axis sections. The WGA images and Φ colormaps for the long-axis sections show that the cardiomyocytes at the edges of the ventricular wall are aligned parallel to the viewing plane (Figure. 3, Supplementary Figure. 12). In the Φ colormaps we show this layer in yellow tones, and it is evident that it extends all the way to the apex (Figure. 3, Supplementary Figures 11 and 12). A complex geometry emerges near the apex region, where the bands of longitudinal myofibers extend, intertwine, and continue into the opposite ends of the ventricular walls (Figure. 3 and Supplementary Figure. 13 a-d).

**Figure. 3:**
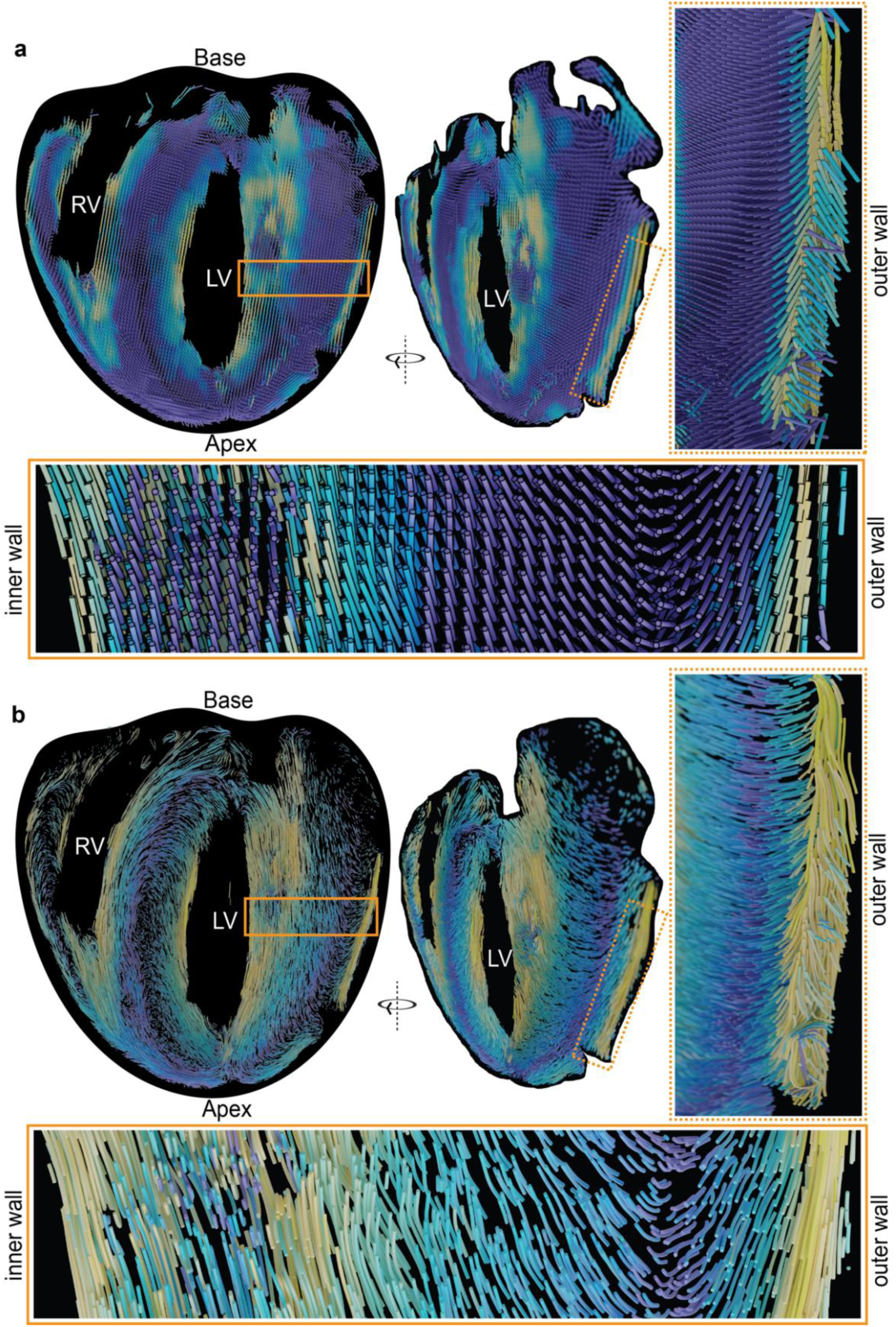
Evidence for longitudinal fibers from the analysis of a long-axis section. Estimated myofiber orientations are shown as glyphs (**a**) and streamlines (**b**) for the mouse LA sections. In each panel, a 3D visualization of the orientations is shown using either glyphs or streamlines, with a view obtained by rotation in a clockwise direction as shown on the right (Methods). The colors follow a parula colormap, where the blue and yellow tones indicate cell orientations that are in and out of the short-axis plane, respectively. These visualizations reveal the continuity of cell orientations across the entire long-axis section, from apex to base.

In order to visualize the continuation of longitudinal fibers at the apex, we extended our sectioning towards the apex region along the short-axis plane (Supplementary Figure. 15) (Methods). Here the θ angle, depicting the component of cardiomyocyte orientation in the XY plane, shows a smooth transition, similar to that in the mid-ventricular short-axis sections (Figure. 2). The Φ angle reveals three longitudinal fiber systems forming a confluence (Figure. 4 and Supplementary Figure. 14). At the apex, these longitudinal fibers turn upwards towards the base of the heart and then continue to the adjacent boundaries of the heart wall. Those from the left ventricular outer wall are connected to the right ventricle outer wall and inner chamber walls (Figure. 4, Supplementary Figures 11 and 13). Thus, the longitudinal myofiber system forms a continuum from the ventricles to the apex (Supplementary Figure. 15), one that is structurally distinct from the circumferential system. To conclude, we have discovered a new prominent fiber system which runs in the long axis direction and is a conserved feature in the ventricular walls of four-chambered hearts.

**Figure. 4:**
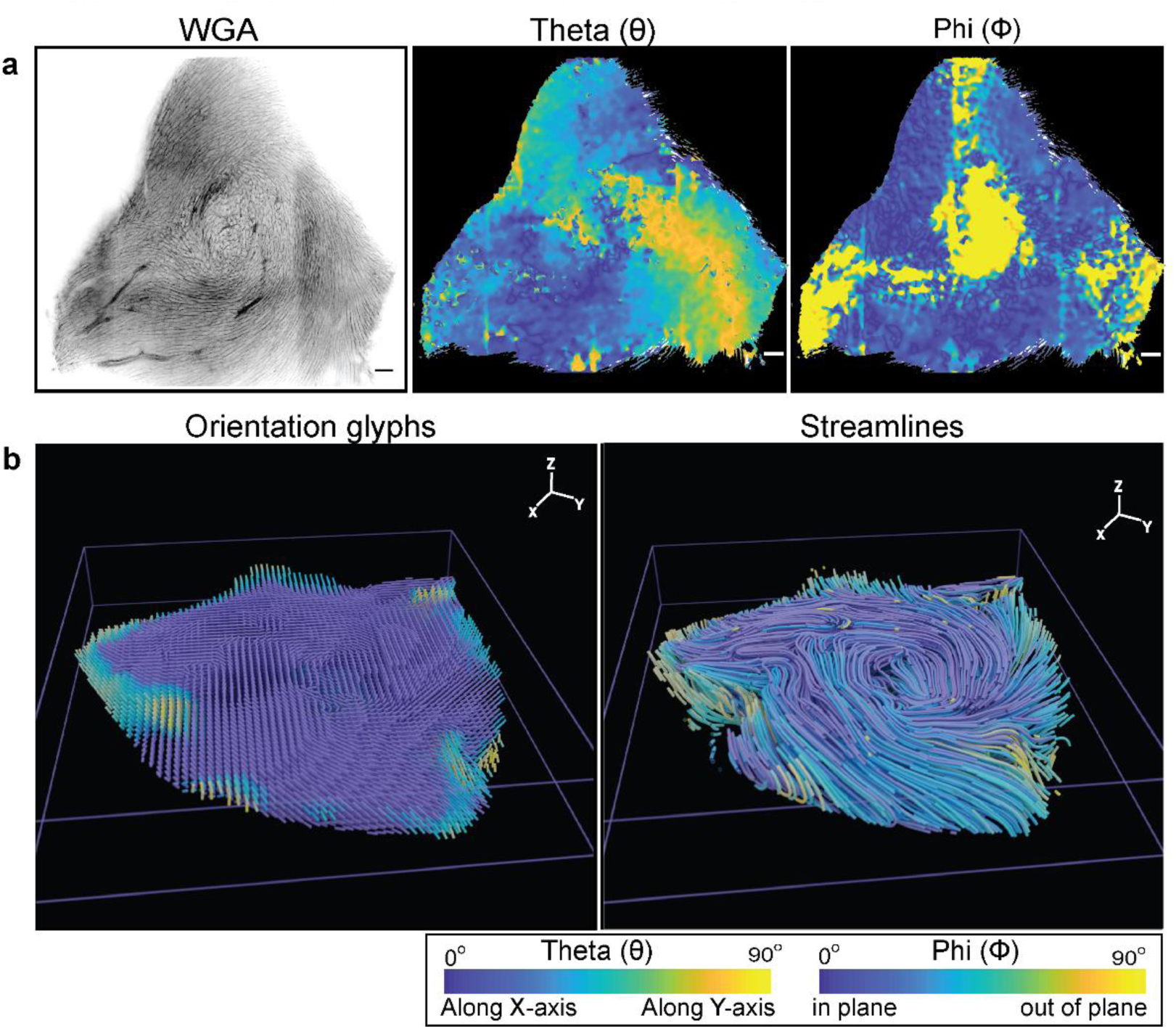
Analysis of a short-axis section from the apex region. **a.** WGA in grayscale, and θ and Φ colormaps of the representative apex section. The colormap scale is as indicated, with the scale bar being 100 microns. **b.** Orientation glyphs and streamlines for the apex section shown in a.

**Figure. 5:**
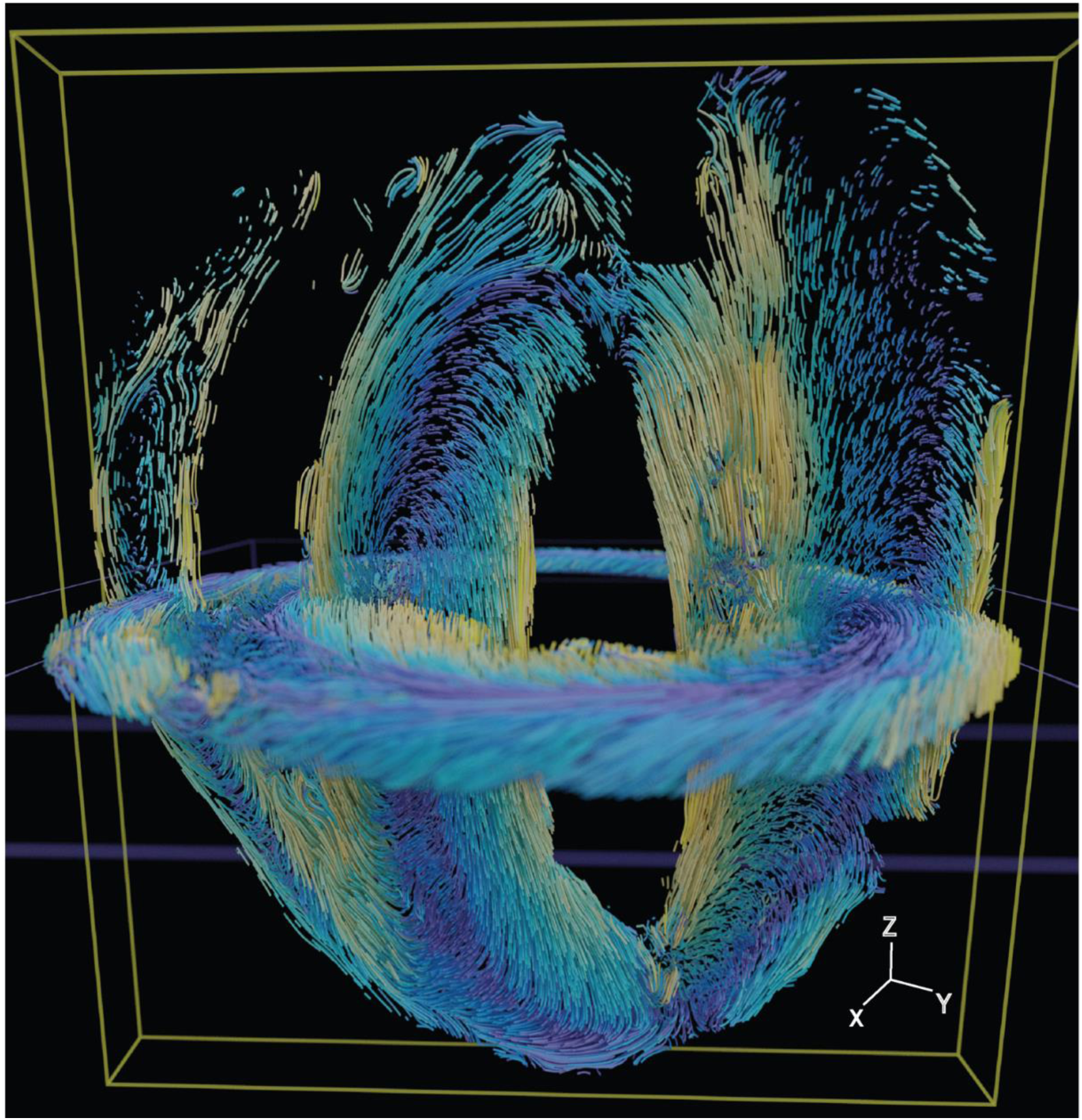
A model for orthogonal myofibers in heart ventricular walls. A model for the longitudinal and circumferential myofibers in the heart ventricle walls, obtained by superposition of the reconstructions of the short-axis and long-axis sections from different mouse hearts. The structure tensor based orientations are visualized as streamlines (Methods). The colors follow a parula colormap, where the blue and yellow tones indicate orientations in or out of the short-axis plane, respectively.

## Discussion

Current heart wall contraction and electrical wave conduction models have only considered the circumferential layers of myofibers in the heart wall (*3, 4*). Our discovery of a new distinct longitudinal fiber system in the ventricular walls of four-chambered hearts could lead to a major shift in our understanding of heart function. During heart contraction, the ventricular walls undergo a wringing motion from the apex towards the base, accompanied by a slight shortening of the heart along its long-axis direction. Together these motions produce sufficient ejection force to empty the chambers. While the bulk of the contraction is carried out by the circumferential myocardial fibers, the continuum of longitudinal fibers might play a significant role in its long-axis shortening. Additionally, we have observed that the four bands of longitudinal fibers continue upwards to the base from the apex (Supplementary Figure. 13) and are anchored to the atrioventricular valves (Supplementary Figure. 16). The opening and closing of the atrioventricular valves must be synchronized with the contraction of the left/right chambers, but currently, no model exists that connects myofibers to the atrioventricular valves. Since the longitudinal fibers are anchored to the valves, we postulate that their opening and closing might partly be controlled by the contraction and relaxation of the longitudinal myofiber system.

In addition to their mechanical function, the longitudinal myofibers might play a complementary role in facilitating electrical conduction in the heart wall. The electrical signals from the sinoatrial node propagate to the atrioventricular node to cause the ventricles to pump blood, via a network of Purkinje fibers (*24, 25*). Analyses have shown that the circumferential fibers, together with their associated transmural turning from the outer wall to the inner wall, can explain the point-to-point time to arrival maps under a model of anisotropic diffusion (*4*). Moreover, the transmural turning has been shown to play a role in minimizing diffusion bias and mitigating the potentially harmful effects of stochastic propagation (*3*). However, how the timing of the contraction is controlled at any short-axis section is an open question. We hypothesize that the propagation of the conduction wave through the heart wall along the transmural direction is mediated by the circumferential fibers (*3, 4*) and that the longitudinal fibers facilitate the propagation of the signal from the apex to the base in a vertical direction. This would then explain the overall timing behavior observed in the contraction of the ventricles and the need for these two orthogonal fiber systems.

Orthogonal fiber systems are, in fact, not restricted to mammalian hearts and similar fiber organizations have been reported in other organs and entire animal body plans (*26–28*). At the organ level, the co-existence of a circumferential and longitudinal fiber system is known to be important for the smooth muscle peristaltic motions seen in esophageal and intestinal tubes (*26, 27*). In planarians, in addition to the circumferential and longitudinal muscle fibers, a diagonal fiber system has been reported (*28*). However, a crucial difference between the heart and other systems is that in the latter, the orthogonal arrays occupy an equal volume (*26–28*). In contrast, our reconstructions show that the longitudinal myofiber systems account for only a small fraction of the myofibers in the heart wall, suggesting that they have highly specialized roles. It remains to be seen whether perturbations of the longitudinal fiber systems are associated with heart diseases including cardiomyopathies, myocardial infarction or electrical conduction disorders. Moreover, the embryonic stage of heart development at which the orthogonal longitudinal myofiber array begins to form in the heart wall is not known. The myofiber geometry at the micron scale reconstructed in our study lays a foundation for examining these questions and for studying the association between pathological heart conditions or disorders and fiber organization.

## Materials and Methods

### Tissue clearing

The mouse and rat hearts were collected from wild-type female C57BL/6 and Wistar strains, respectively. All heart samples were perfused during the collection with heparinised 1X PBS to remove blood clots, followed by 4% paraformaldehyde (PFA). The fixed hearts were stored in 4° Celsius until further use. To clear the heart tissue we applied the CLARITY method (*29*) with the following modifications. The fixed mouse hearts were transferred to hydrogel monomer solution (PBS, 4% acrylamide, 4% PFA, 0.5% Bisacrylamide and 0.25% photo-initiator 2, 20-Azobis[2-(2-imidazolin-2-yl) propane] dihydrochloride (VA-044, Wako Chemicals USA) for 7 days. For initiating the hydrogel hybridization and polymerization the processed heart tissues were incubated for 3 hours at 37° Celsius. After polymerization, the excess gel material was removed. The tissue was transferred to 50 ml tubes and washed 3 times with PBS, then incubated with a clearing buffer (8% SDS and 4% boric acid in 1X PBS (pH 8.5)) for 20-30 days at 37° Celsius, in a shaking incubator at180 rpm, with the exchange of a new clearing buffer every week. This CLARITY based approach applied to the heart tissue samples resulted in a transparent tissue (Figure. 1A and Supplementary Figure. 1A).

### Sample preparation and imaging

The cleared mouse hearts and uncleared fixed mouse and rat hearts were subjected to short- and long-axis sectioning using a Compresstome VF-300, as illustrated in Supplementary Figure. 1B. The sectioned heart tissues were processed for staining by washing them with PBS 3 times, followed by the application of PBST (PBS + 1% Triton X-100) for 24 hours at 37° Celsius. Subsequently, the heart sections were incubated in 150μg/ml of Alexa Fluor^TM^ 633 conjugated wheat germ agglutinin (WGA, W21404, ThermoFisher) for 24 hours and washed with PBS, 3 times, for 10 minutes each time. The WGA stained cleared and uncleared heart tissues were then transferred into RIMS imaging media (88% Histodenz Sigma D2158 in 20mM Phosphate buffer pH 7.5). The heart tissues were then mounted with fresh RIMS, sandwiched between two coverslips using 500 micron spacers (IS002, SUNjin Lab, Taiwan). The confocal images were acquired using an Olympus FV3000 microscope with an Olympus PlanApo 1.25X and Olympus UCPLFN 20X CorrM32 85mm scale air objective (NA=0.73). For each section, using the 1.25X objective, a lower magnification image encompassing the whole area of the section was obtained, which was then used to map the fields of view using the Olympus fluoView^TM^ software.

Micron scale imaging was carried out using a 20X objective, with each field of view covering 320×320 pixels with an isometric voxel size of 1.98µm^3^. For the cleared and uncleared heart tissues a maximum depth of 300 and 50 µm of image stacks were acquired, respectively. The samples were excited using a 640nm laser line and emission was detected over a range of 650-670nm using high sensitivity spectral detectors (gallium arsenide phosphide photo-multiplier tube).

### Pre-processing of heart tissue image stacks

At the deeper end of the image stacks we observed a poor signal to noise ratio. In order to improve the fluorescence signal, we performed deconvolution and denoising using custom built algorithms. For the deconvolution, we corrected the non-ideal PSF using iterative Richardson-Lucy deconvolution (30) with Total variation regularization. The deconvolved image stacks were then subjected to unsupervised denoising using a dictionary learning method (31). The dictionary was extracted from the shallow layers of image stacks under the assumption that the deeper and shallow layers contained common substructure elements of cells. This set of learned dictionary patches (a sparse 256 element 2D dictionary of patches size 16 x 16) from cardiac tissue was used to computationally clear the noise and improve the visibility of cell structures at the deeper end of the stacks. A separate dictionary file was generated for each individual field of view to prevent structures from one region from influencing the reconstruction in other regions of the heart tissue sample. The deconvolved and denoised image stacks were stitched together to yield a micron scale complete short- and long-axis section, with a depth of about 300 µm. To achieve this, during imaging each individual field view was set to have at least a 25% overlap with its neighboring fields of view. The overlapping fields of view were tiled using phase correlation between adjacent fields of view in the Fourier domain (32). The tiled reconstructions were globally aligned with AHA sectors in a manual fashion, using features including capillary vessels and papillary muscle placement and orientation.

### Cell orientation estimation using a structure tensor

The intensity of the WGA stain, which is absorbed by the cell membranes of the myocytes, was used to estimate myocyte orientation. At each voxel in the deconvolved and denoised image stack, we computed the structure tensor (*33*). We then associated to each voxel the orientation of the direction in which the intensity varied the least, capturing the long-axis orientation of myocytes, by selecting the eigenvector of the structure tensor corresponding to the eigenvalue with smallest magnitude. The mapping of myocyte orientation to coarser spatial scales was carried out by element-wise averaging of local structure tensors, followed by eigenvector decomposition of the smoothed tensors. The cell orientation was then interpreted in terms of its projection in the short-axis plane θ, the component out of the short-axis plane Φ, and the angle αΗ between its projection onto the heart wall tangent plane and the long-axis direction (Figure. 2A).

### Colormap, Glyphs and Streamline generation

The orientation field was represented by a 3-dimensional array of rotated cylinders, referred to as orientation glyphs. The input vector field was approximated by a 3-dimensional grid of equally spaced vertices, down-sampled such that the number of vertices was not larger than 75,000. At each vertex of the down-sampled grid, a cylinder primitive shape was created, and rotated proportionally to the components of the vector field at the vertex position. A parula colormap was applied to the cylinders in a manner that was proportional to the Φ angle, using the arc cosine of the absolute value of the z component of the vector field. Bidirectional streamlines were represented as curves extruded from polylines, whose points were computed as follows. A set of up to 25,000 points were selected from a random sample of voxels from the vector field, and each sample voxel location was the initial point for a streamline. Since the vector field represents the orientation of the tissue, each starting point initialized both a positive and negative streamline, each of which was grown by iteratively adding new points along the polyline. At each new position, the value of the field at that voxel acted to determine the position of the following point along the polyline. A parula colormap was applied to the streamlines in proportion to the Phi angle at the starting point within the vector field. To avoid artefacts arising where the streamlines extended beyond imaged data, a binary mask of the tissue volume with the same dimensions as the vector field served as a boundary condition.

A detailed account of the methods is in Supplementary Information

## Acknowledgements

We are grateful to the Central Imaging and Flow Facility (CIFF) and the Animal Care and Resource Center (ACRC) at the Bangalore Life Science Cluster, India. The ACRC was partially supported by the National Mouse Research Resource grant (BT/PR5981/MED/31/181/2012;2013-2016 & 102/IFD/SAN/5003/2017-2018) from the Department of Biotechnology, India. M.S acknowledges funding support from inStem core grants from the Department of Biotechnology, India, DBT/Wellcome Trust India Alliance Intermediate Fellowship (IA/I/14/2/501533), EMBO Young Investigator Programme award, CEFIPRA (5703–1) from the Department of Science and Technology, SERB-EMR grant (CRG/2019/003246) and DBT-BIRAC (BT/PR40389/COT/142/6/2020) grant. P.S.D is supported by the DBT/Wellcome Trust India Alliance Intermediate Fellowship (IA/I/16/1/502367), Rajiv Gandhi University of Health Sciences (RGUHS), Scientist Development Grant (15SDG23250005) from American Heart Association (AHA), Department of Science and Technology (DST/CRG/2019/005401) and inStem core funding. K.S. is grateful to the Natural Sciences and Engineering Research Council of Canada (NSERC) and to the Fonds de Recherche du Québec Nature et Technologies (FRQNT) for research funding. D.D is supported by a CSIR-SRF.

## Author Contribution

K.S. and M.S. conceived of and supervised the project. D.D. lead the biological aspects of the project, including tissue preparation, confocal imaging and data analysis, in consultation with M.S. T.A.S. lead the computational aspects of the project, including the development of the pipeline for reconstruction, the implementations of algorithms, and the analysis of orientation data, in consultation with K.S. T.F.W.S. developed the high-resolution visualizations of streamlines and glyphs used in some of the figures. P.S.D. provided guidance with the animal studies. D.D., T.A.S, K.S. and M.S. interpreted the results and wrote the manuscript.

---

**Supplementary Figure. 1:**
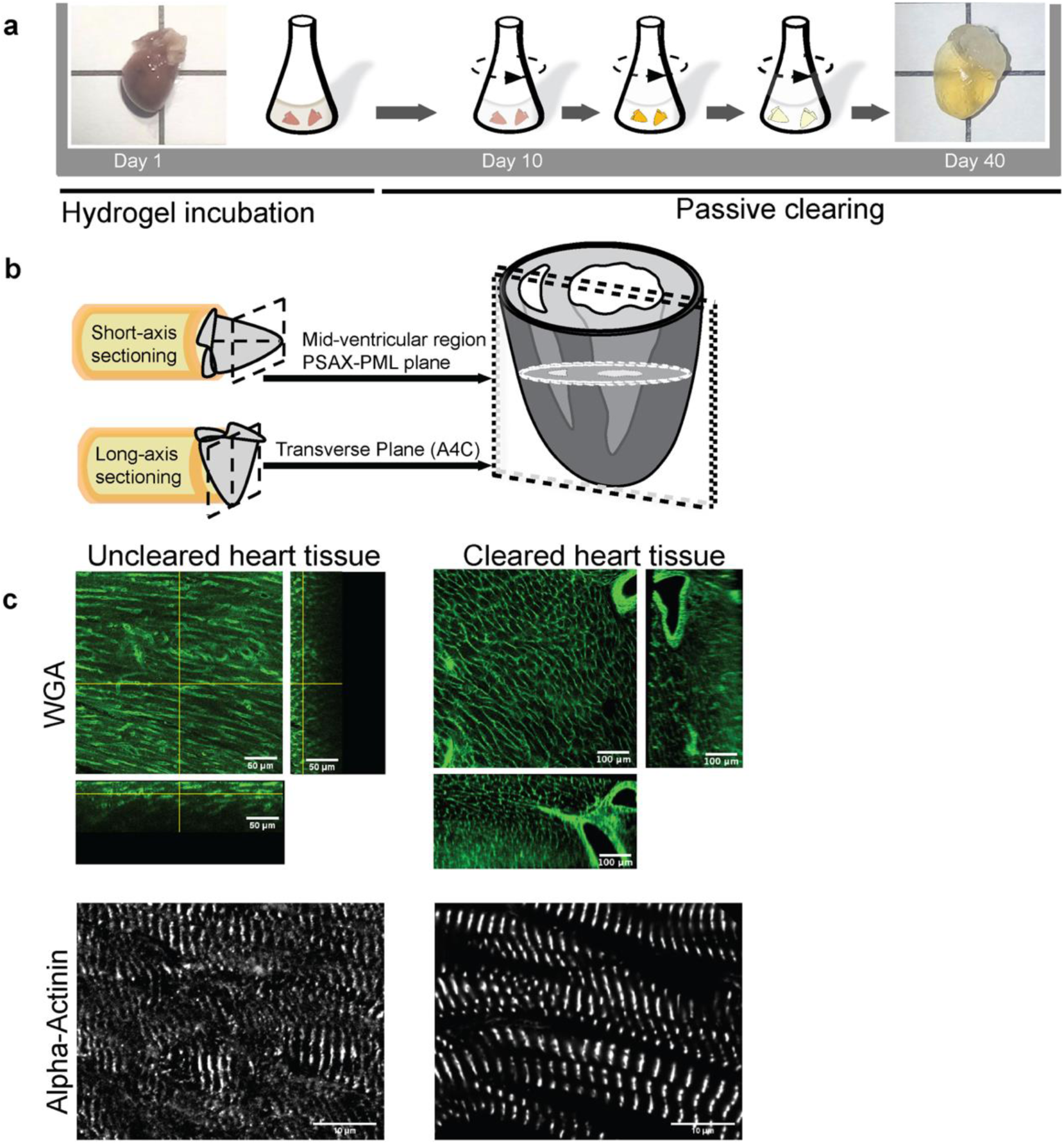
Mouse heart tissue preparation for imaging. **a.** A schematic illustration of the Clarity method. The harvested heart tissue is incubated in a hydrogel/PFA mixture at 4° Celsius for a minimum of 9 days, following which it is stored until further use. The solution mixture with the heart tissue is then polymerized at 37° Celsius. The heart tissue is then excised from the polymerized hydrogel and shaken at 37° Celsius with a clearing solution until it attains a desirable transparency (See Methods for details). **b.**The clarified heart is sectioned along its Short-axis or Long-axis. For the short-axis, a mid-ventricular region that approximates the PSAX-PML (parasternal short-axis – papillary muscle level) plane was chosen. For the Long-axis, a transverse plane that represents the A4C (apical four chambers) plane was used. **c.** A comparison of uncleared and cleared heart tissue sections stained with WGA and the alpha-actinin antibody. The scale bar is 50 and 100 microns for WGA uncleared and cleared tissue images, respectively. The scale bars for the alpha-actinin images are 10 microns.

**Supplementary Figure. 2:**
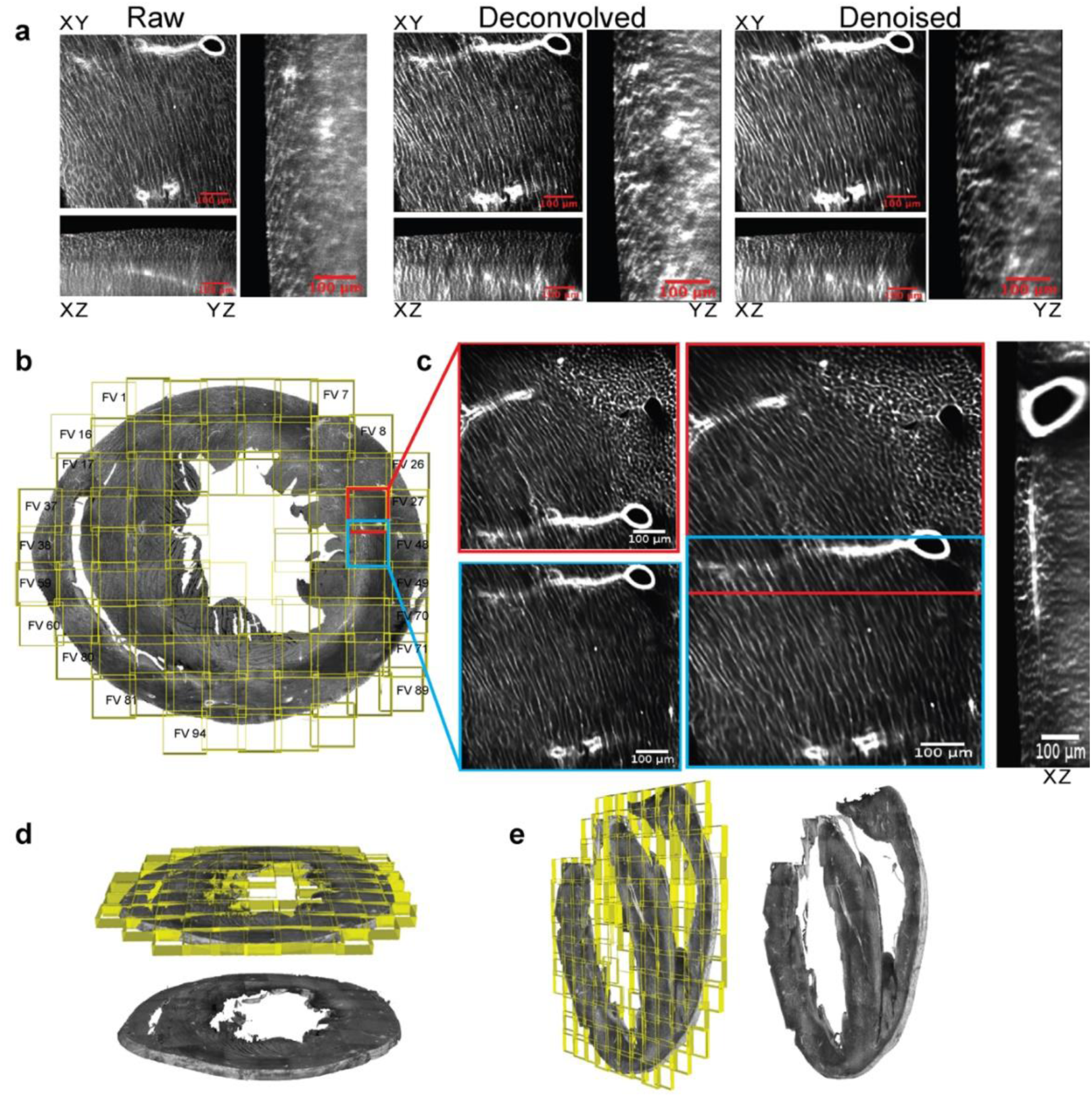
Preprocessing and stitching individual fields of view. **a.** A representative field of view (left), followed by deconvolution (middle) and denoising (right). The orthogonal views (XZ and YZ) show an improvement in the signal to noise ratio towards the deeper Z-sections. The scale bar is 100 microns. **b.** Individual fields of view (FVs) overlayed as a grid on the reconstructed full short-axis section. The individual FVs were imaged using a snake pattern, row by row. Each FV is 320 microns^2^ in dimension and has a 25% overlap with its neighboring FVs. **c.** Left panel; An example showing the stitching of two neighboring FVs (_FV_28 and _FV_47, shown in red and blue boxes, respectively). Middle panel; A zoomed in view of the stitched result, showing the alignment of features in the common region. Right panel; An XZ/YZ view of the common region. The scale bar is as indicated. **d-e.** 3D-views of the stitched short-axis and long-axis sections, with the individual FVs shown in yellow in an overlayed grid pattern.

**Supplementary Figure. 3:**
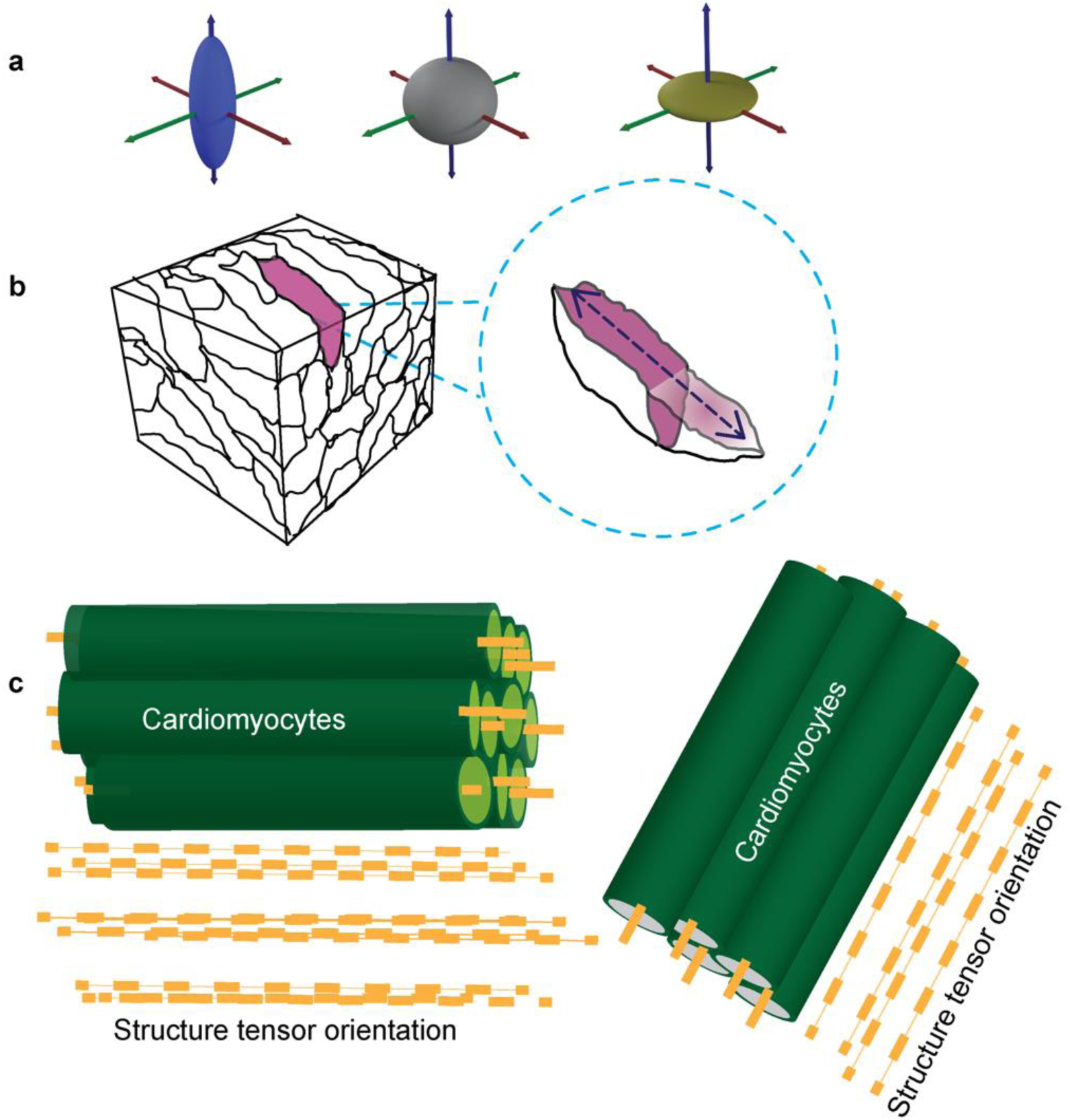
The structure tensor method for orientation estimation. **a.** Sample structure tensors. Left: an elongated tensor (the preferred orientation is in the direction of elongation) Middle: an isotropic tensor (with no preferred local orientation). Right: a plate-like structure (with two preferred orientations). **b.** Illustration of a block of heart tissue in which a single cardiomyocyte is highlighted in purple and is described by an ellipsoidal tensor, whose major axis is along the long-axis of the cardiomyocyte. **c.** Illustration of cell orientation estimation using a structure tensor method (Methods). The structure tensor eigenvectors with the smallest eigenvalue (yellow dashed lines) align to the long-axis direction of the cardiomyocytes (green cylinders).

**Supplementary Figure. 4:**
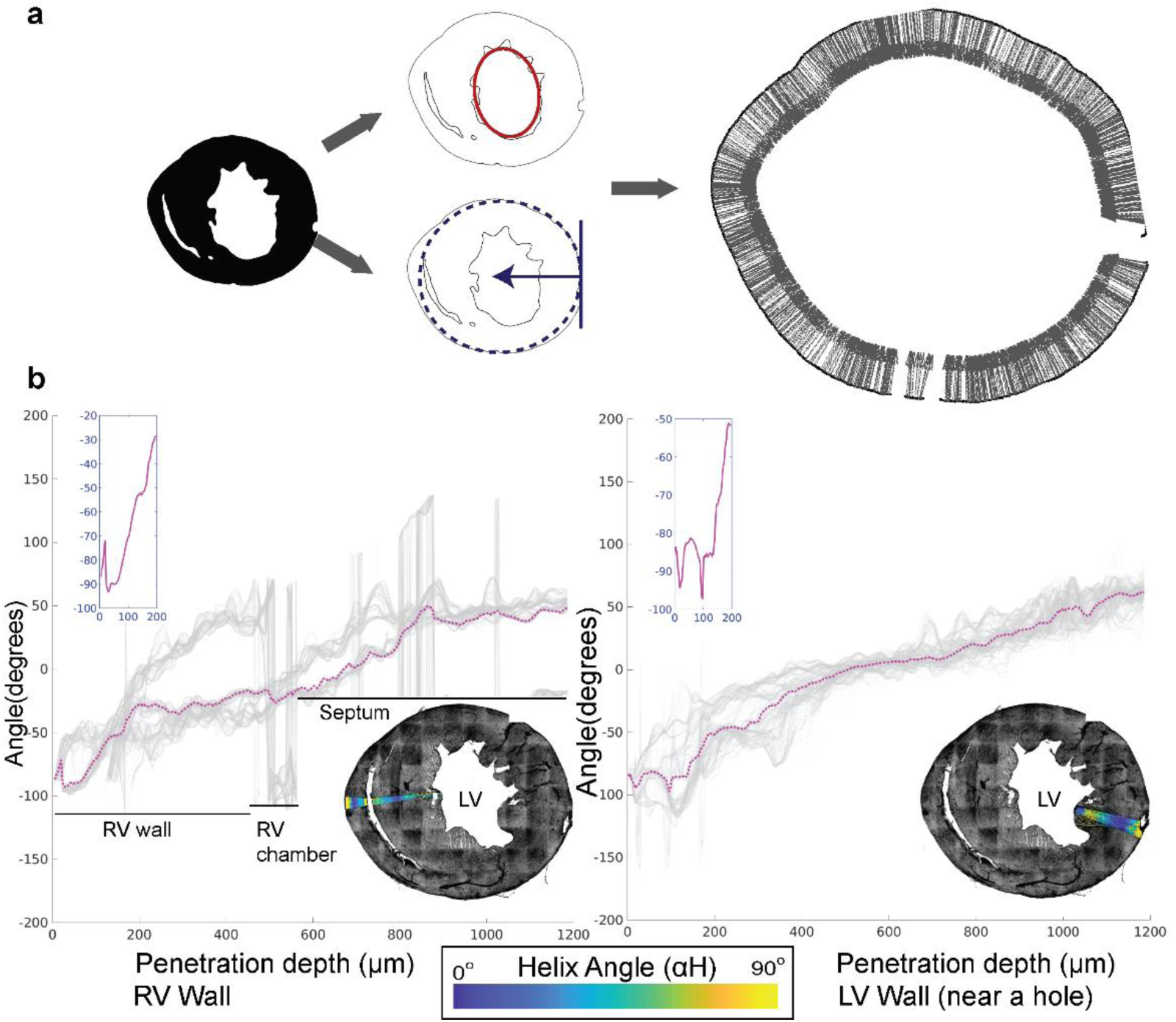
Estimation of the helix angle from short-axis sections. **a.** Helix angle calculation: Masking followed by centroid estimation (top); Masking the short-axis section and estimating the tangent plane and normal for the penetration axis (bottom). **b.** Average helix angle plots from outer to inner wall of the right (left side) and left (right side) ventricular walls. The individual pie shaped sectors that were averaged are overlayed using a parula colormap on a maximum intensity Z-projection of the WGA-stained short-axis section shown in grayscale (middle right).

**Supplementary Figure. 5:**
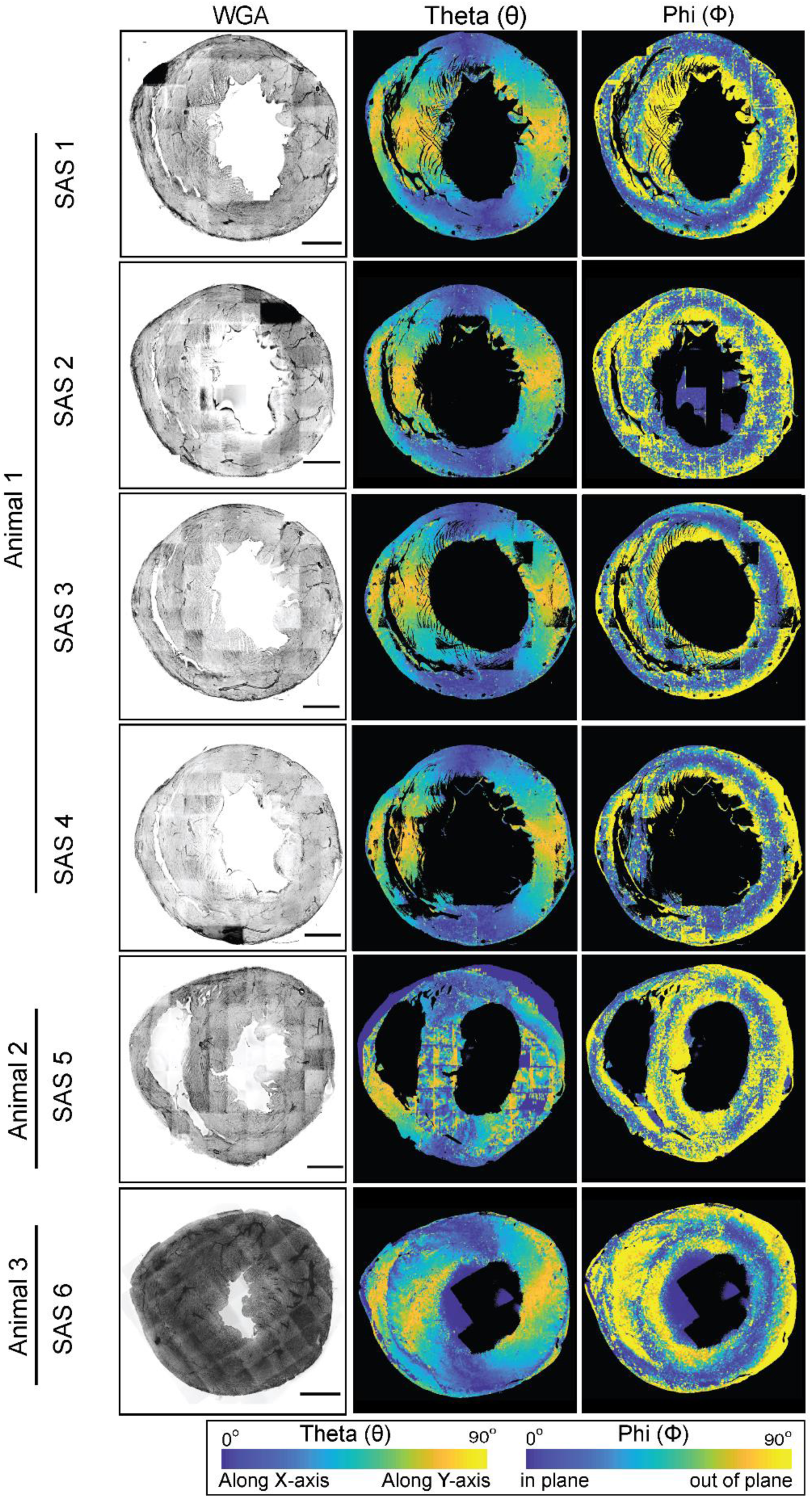
WGA staining and angular colormaps of different short-axis sections. A maximum intensity Z-projection of the WGA-stained short-axis section from a mouse heart is shown in grayscale, with the Φ and θ angles for cell orientations shown using parula colormaps, with the color bars as indicated. The scale bar is 1000 microns.

**Supplementary Figure. 6:**
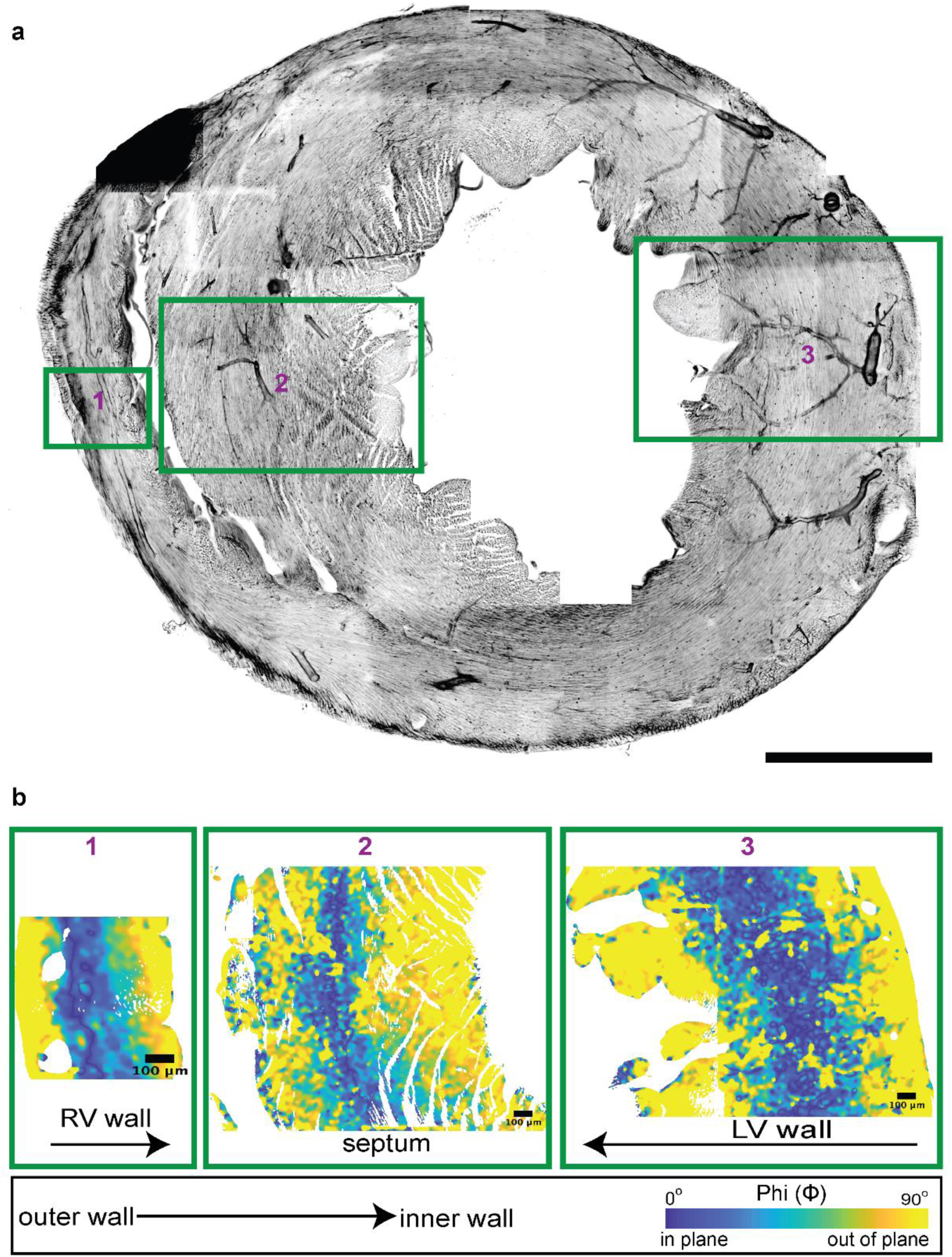
Φ angle colormaps of the inner walls in a short-axis section. **a.** A maximum intensity Z-projection of the WGA-stained short-axis section from a mouse heart is shown in grayscale. **b.**Zoomed in regions of Z-sections of the RV, Septum and LV walls with Φ shown using a parula colormap. The scale bar is 1000 microns and the color bars are as indicated.

**Supplementary Figure. 7:**
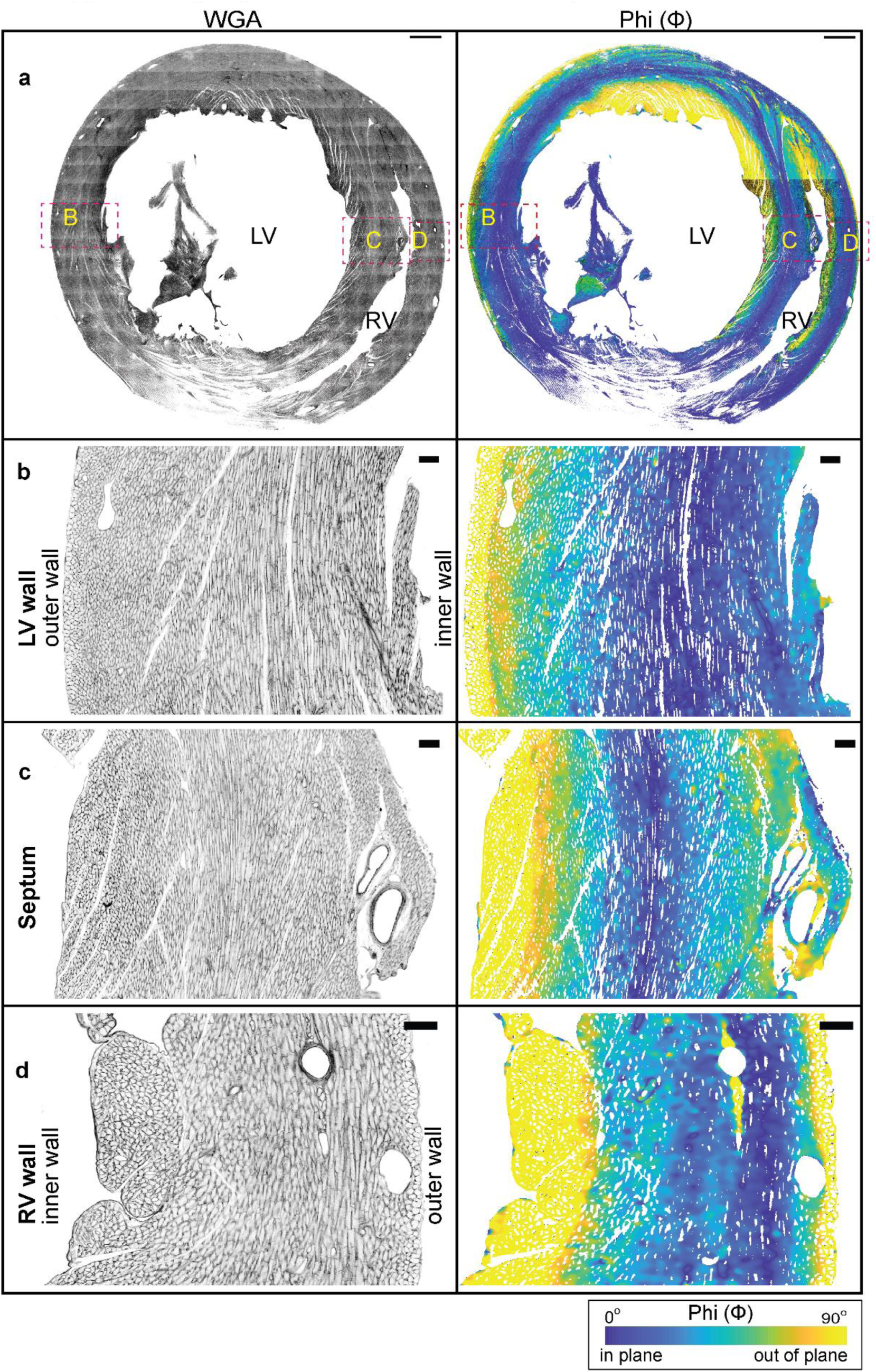
Short-axis section analysis of a rat heart. **a.** A short-axis view of the ventricular chambers of a rat heart, sectioned at PSAX-PML (parasternal short-axis – papillary muscle level). A maximum intensity projection of the WGA stain is shown in grayscale, with the Φ angle shown using a parula colormap. The scale bar is 1000 microns. **b-d.** Zoomed in regions from the left ventricle, septum and right ventricle, with the WGA stain shown in grayscale and the Φ angle shown using a parula colormap. The scale bar is 100 microns and the color bars are as indicated.

**Supplementary Figure. 8:**
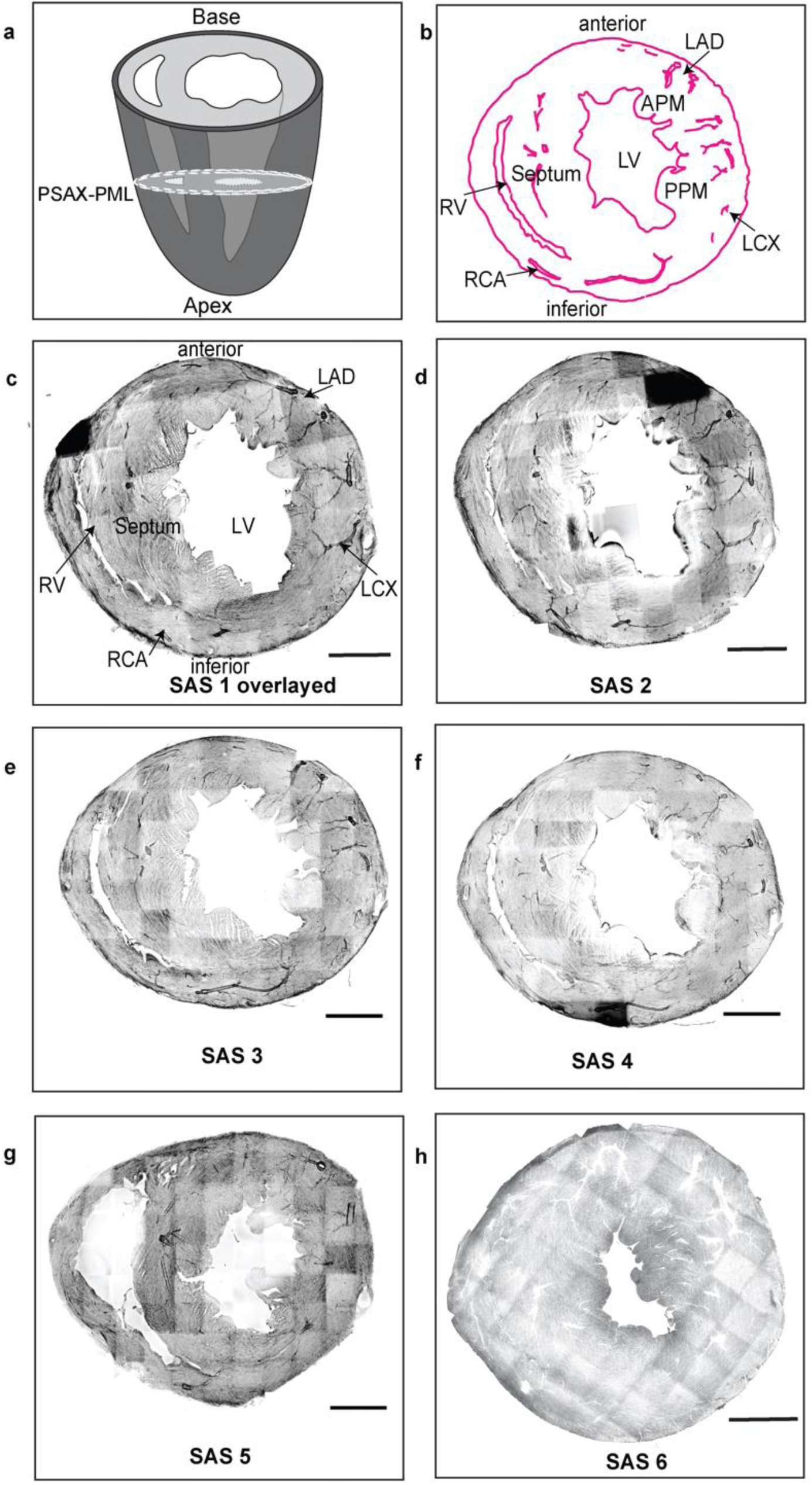
Alignment of different short-axis sections. **a-c.** Boundaries of the short-axis sections (PSAX-PML section) from a WGA maximum intensity Z-projection, with labels as illustrated. Using the major arteries, blood vessels and papillary muscles in the LV chamber, the anterior and inferior orientations were aligned for the different short-axis sections. RCA-Right Coronary Artery, LCX-Left Circumflex Artery, LAD-Left Anterior Descending Artery, APM-Anterior Papillary Muscle, PPM-Posterior Papillary Muscle, RV-Right ventricle, LV-Left Ventricle. **d-h**. The aligned datasets are shown as maximum intensity projections of the WGA stain images. The scale bar is 1000 microns.

**Supplementary Figure. 9:**
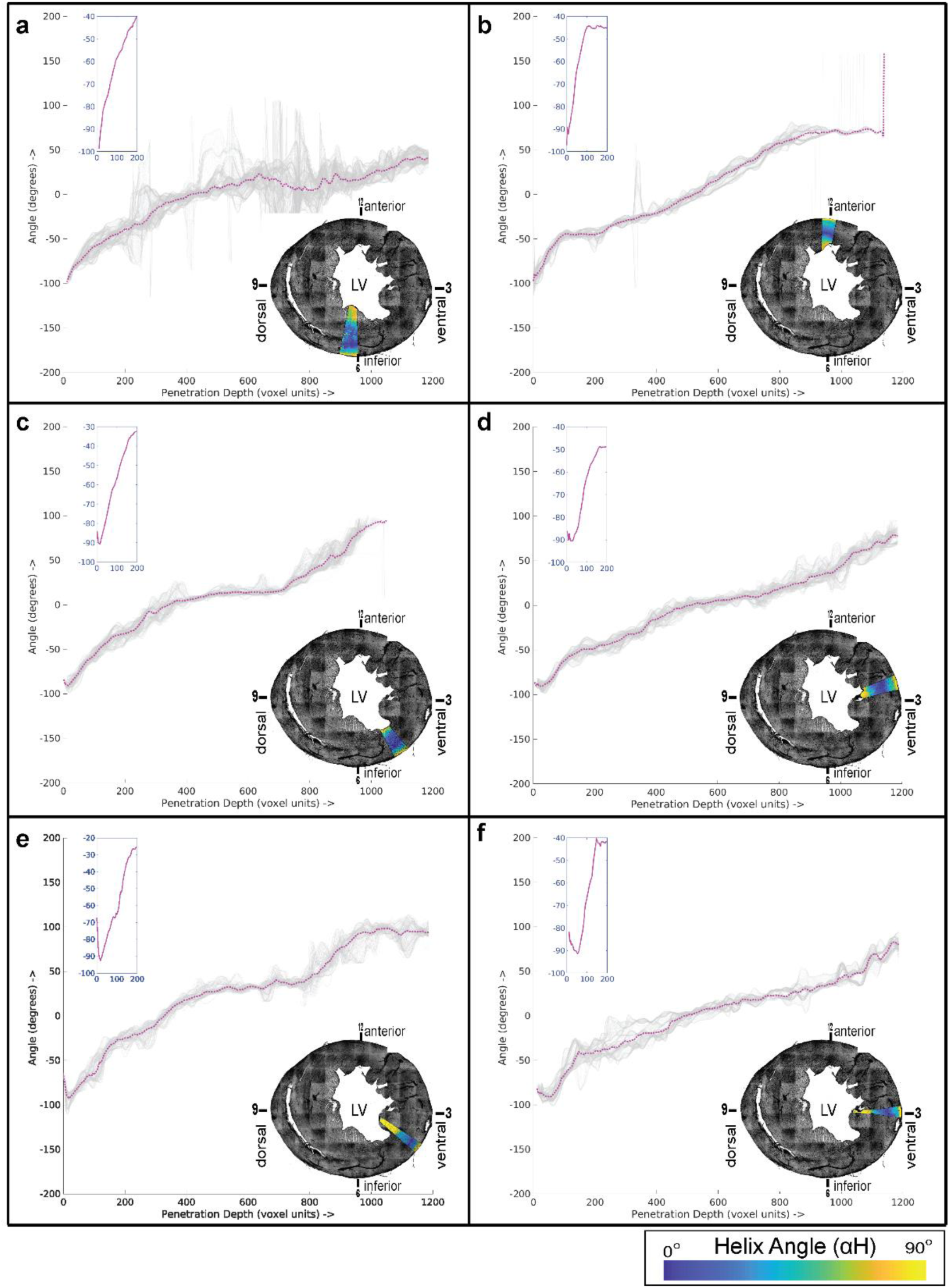
Variation of αΗ from the outer to the inner ventricular walls. An examination of αΗ in a short-axis view of a mouse heart ventricle, sectioned at the PSAX-PML (parasternal short-axis – papillary muscle level), lower right of each panel. The αΗ values of selected pie-shaped sectors* of the LV region from one representative data set of heart sections (SAS3) are plotted as a function of the penetration depth from the outer to the inner walls. Penetration depth is depicted as distance along the X-axis. The mean αΗ is plotted as a dotted magenta line and the αΗ measurements along distinct penetrations are shown in gray. *The values plotted here derive from sectors spanning 10 degrees each, as illustrated.

**Supplementary Figure. 10:**
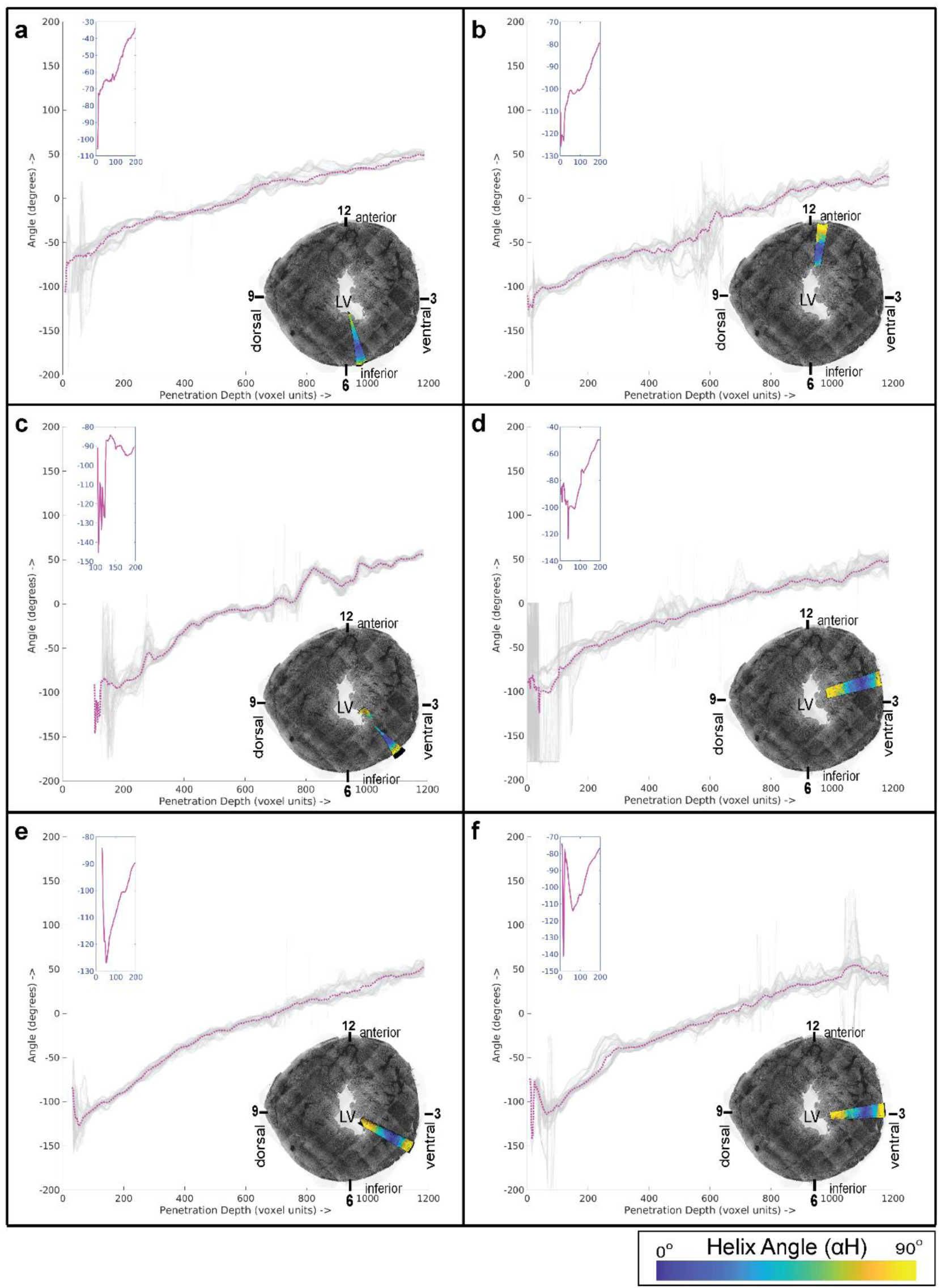
αΗ plots from the subsequent section, dataset SAS5. Custom pie-shaped sectors of the ventricular walls are selected. Each sector spans 10 degrees and is overlayed as a magenta wedge on a maximum intensity Z-projection of the WGA stain at the bottom right of each panel. The αΗ values are plotted along the penetration from the outer to inner walls, depicted using distance in microns along the X-axis. The mean is highlighted as a magenta curve in the plots and a zoomed in version of the outer-ventricular region is provided at the top left of each panel.

**Supplementary Figure. 11:**
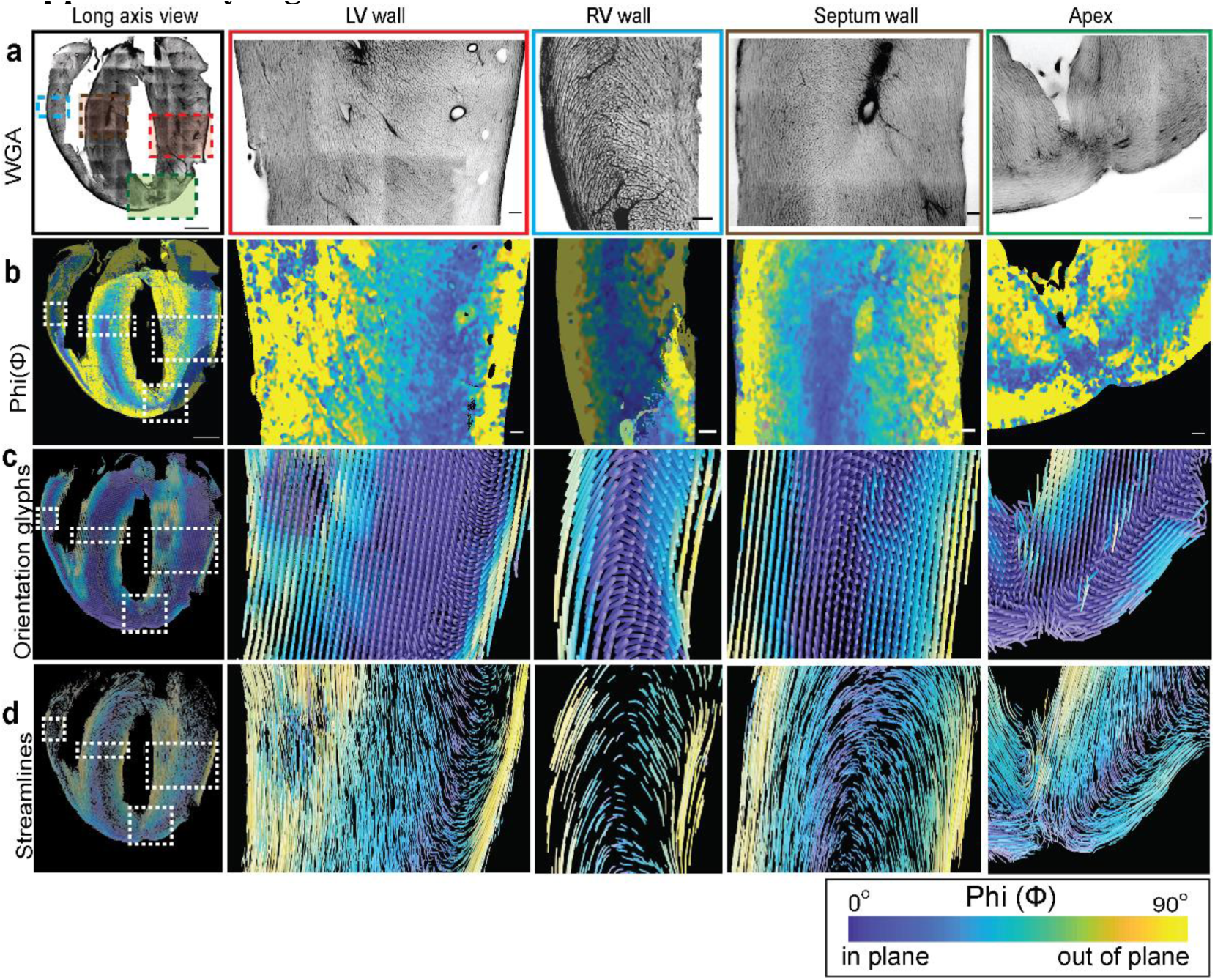
A detailed view of a long-axis section. **a.** WGA stain**. b.** Φ angle colormaps. **c.** Estimated orientations as glyphs. **d.** Estimated orientations as streamlines. The colors follow a parula colormap, where the blue and yellow colors indicate in- and out-of short-axis plane cell orientations, respectively. Magnified views of the left ventricle (2^nd^ column), right ventricle (3^rd^ column), septum wall (4^th^ column) and the apex region (5^th^ column) are shown for the regions in panel A. The color bar for the Φ angle is as indicated. The scale bar for the complete view is 1000 microns and for the zoomed in regions is 100 microns.

**Supplementary Figure. 12:**
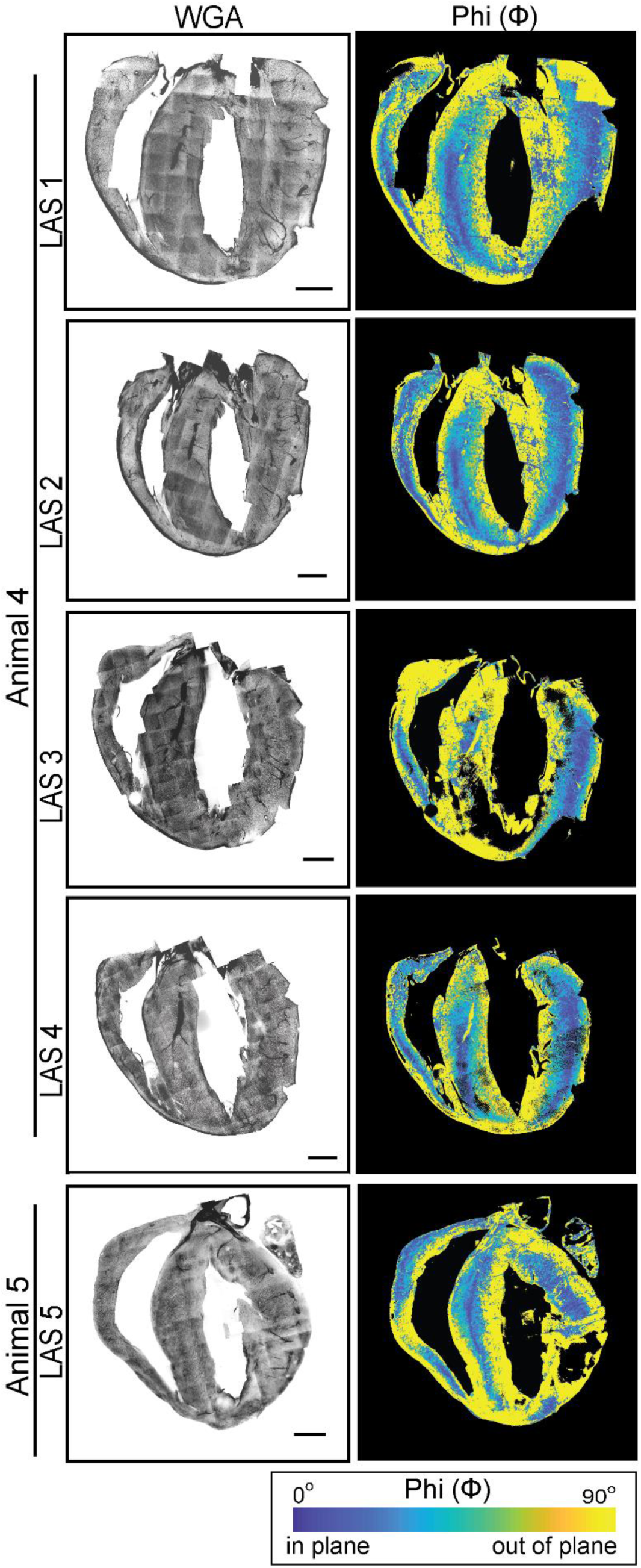
WGA staining and Φ angle colormaps of different long-axis sections. Five mouse heart ventricle long-axis sections are shown as maximum intensity Z-projections of the WGA stain, with the Φ angles of cell orientation shown with a parula colormap. The color bar is as indicated and the scale bar is 1000 microns.

**Supplementary Figure. 13:**
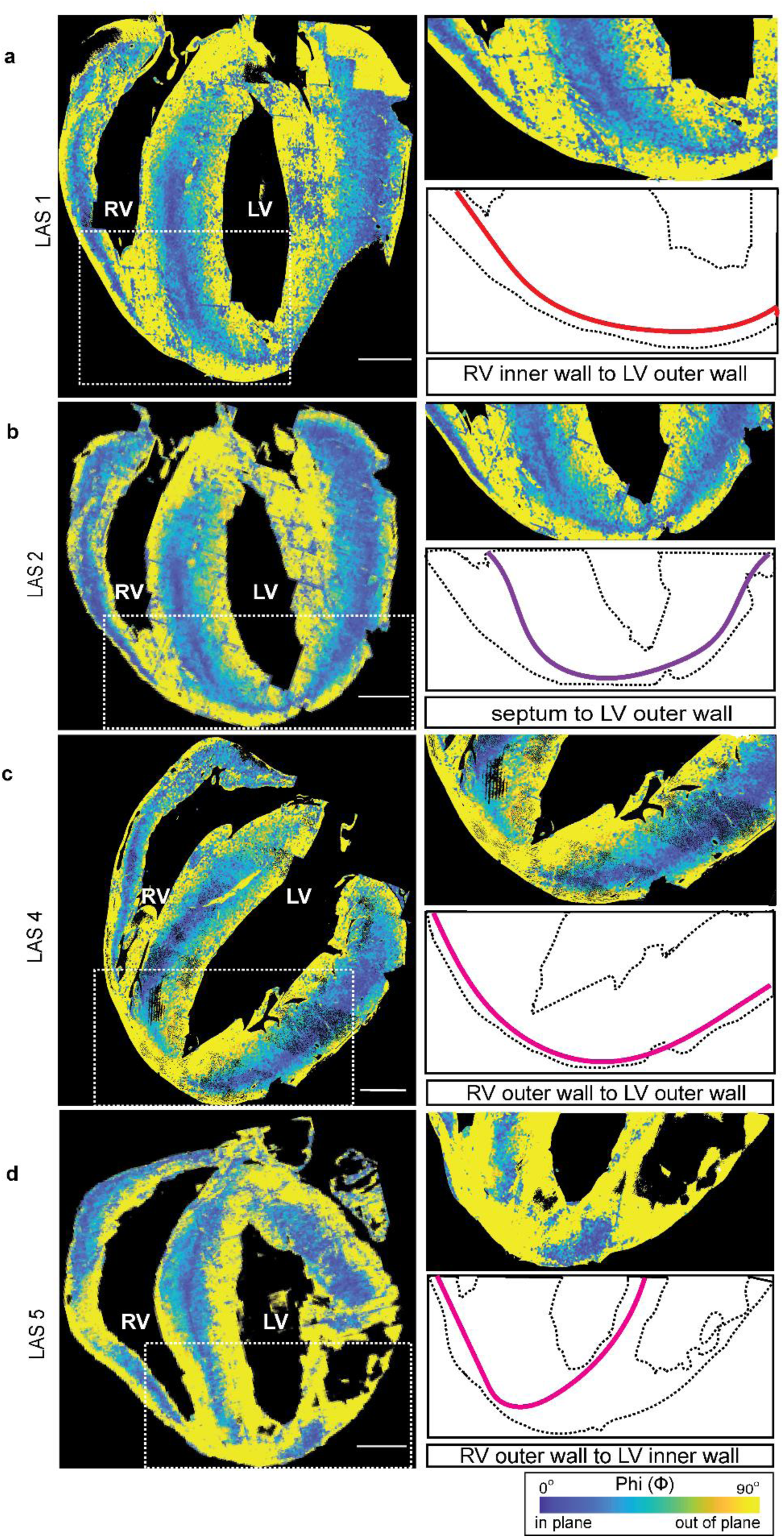
Connections of longitudinal fibers at the apex. Mouse heart ventricle long-axis sections, with Φ angles shown using a parula colormap, with a white dotted box highlighting the apex region under consideration (left column). A zoomed in view of the highlighted apex region from the respective long-axis sections (middle column). An illustration of longitudinal connections between different ventricular walls (right column). The continuity of the longitudinal fibers from the septum to the left ventricle outer wall, from the right ventricle inner wall to the left ventricle outer wall, from the right ventricle outer wall to the left ventricle outer wall and from the right ventricle inner wall to the left ventricle inner wall, is highlighted in **a**, **b**, **c** and **d,** respectively. The scale bar is 1000 microns.

**Supplementary Figure. 14:**
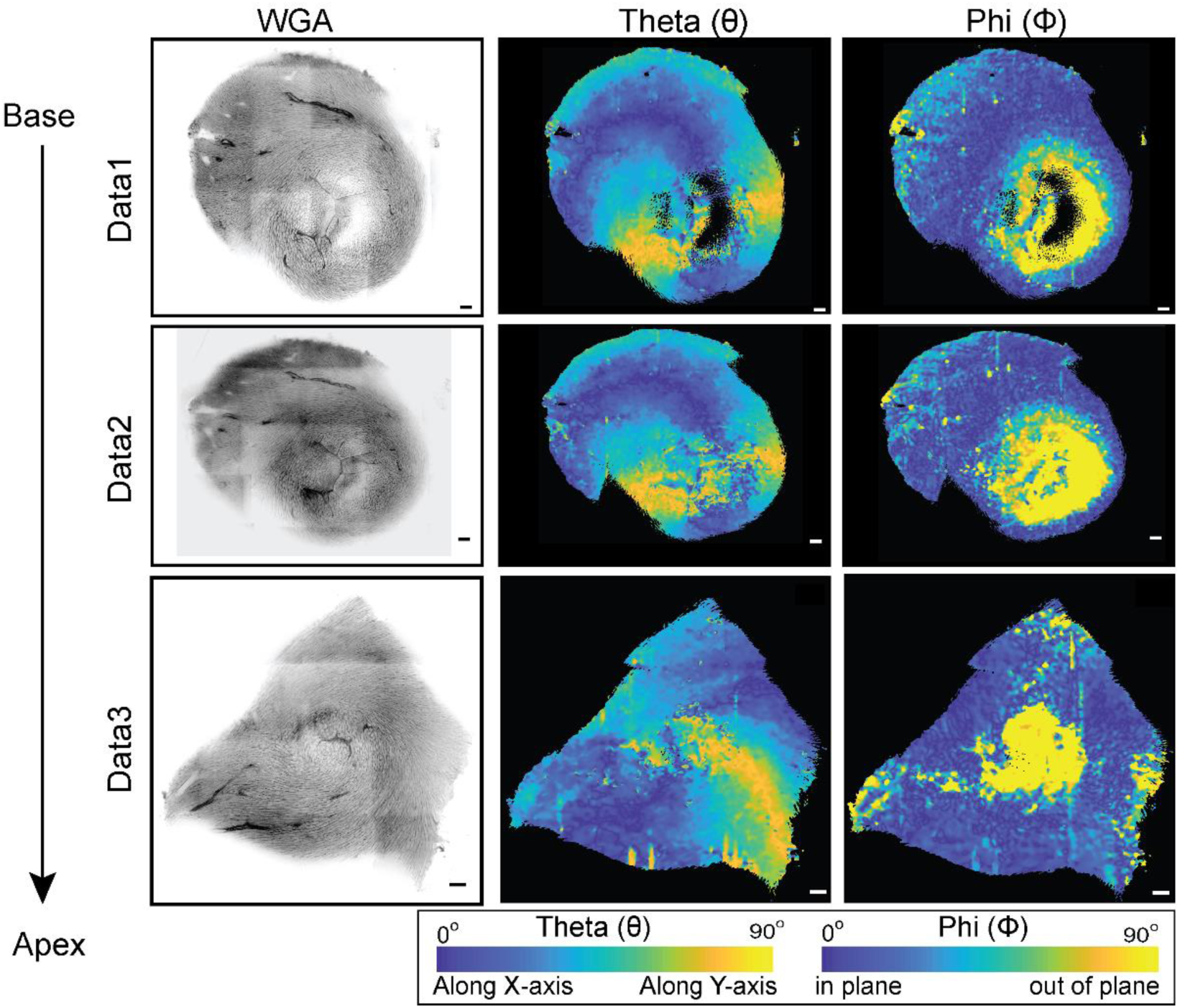
WGA staining and angular colormaps of different apex sections. Three representative Z-planes from four mouse heart ventricle apical serial sections are shown with WGA staining (left), and the θ, and Φ angles related to cell orientation using parula colormaps. The color bars are as indicated and the scale bar is 100 microns.

**Supplementary Figure. 15:**
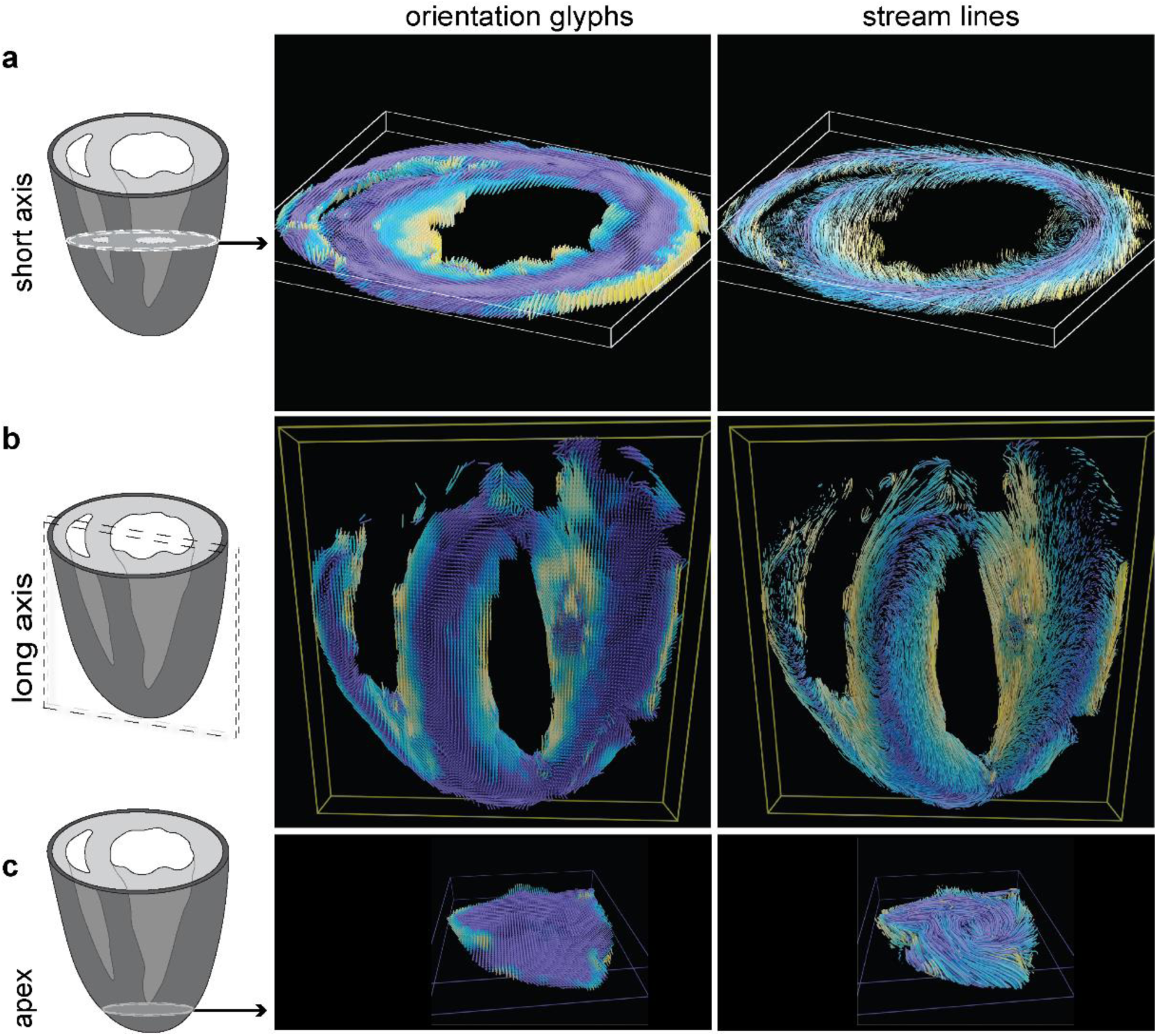
Longitudinal and circumferential myofibers in the heart ventricular walls. Schematics of the heart ventricle walls and the heart sections analyzed (left); **a.** A short-axis (mid-ventricular region) section, **b.** A long-axis (transverse section) and **c.** An apical section. Structure tensor based orientations are visualized as glyphs (middle) or streamlines (right) (Methods). The colors follow a parula colormap, where the blue and yellow tones indicate orientations that are in or out of the short-axis plane, respectively.

**Supplementary Figure. 16:**
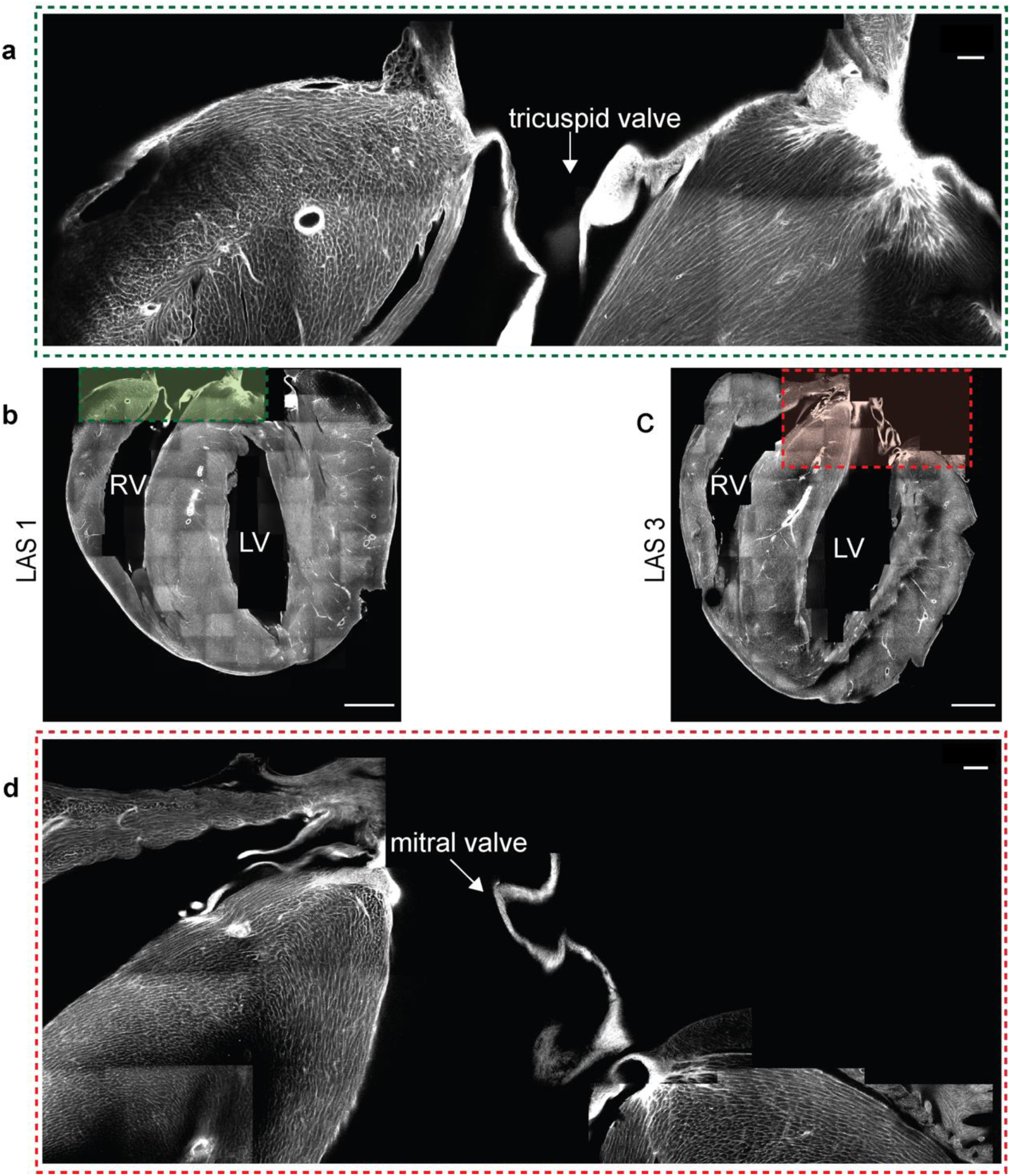
Connections of longitudinal fibers at the atrioventricular valve region. Representative images highlighting the anchor points of the longitudinal fibers of the right ventricle and the septal outer and inner walls (top) and the longitudinal fibers of the left ventricle and the septal outer and inner walls (bottom). **a. and c.** WGA maximum intensity Z-projection of the whole long-axis section. **b. and d.** Zoomed in regions of the right and left AV valve as indicated, WGA (in grey) and streamlines (in parula colormap). The scale bar is 1000 microns.

**Supplementary Table 1:**
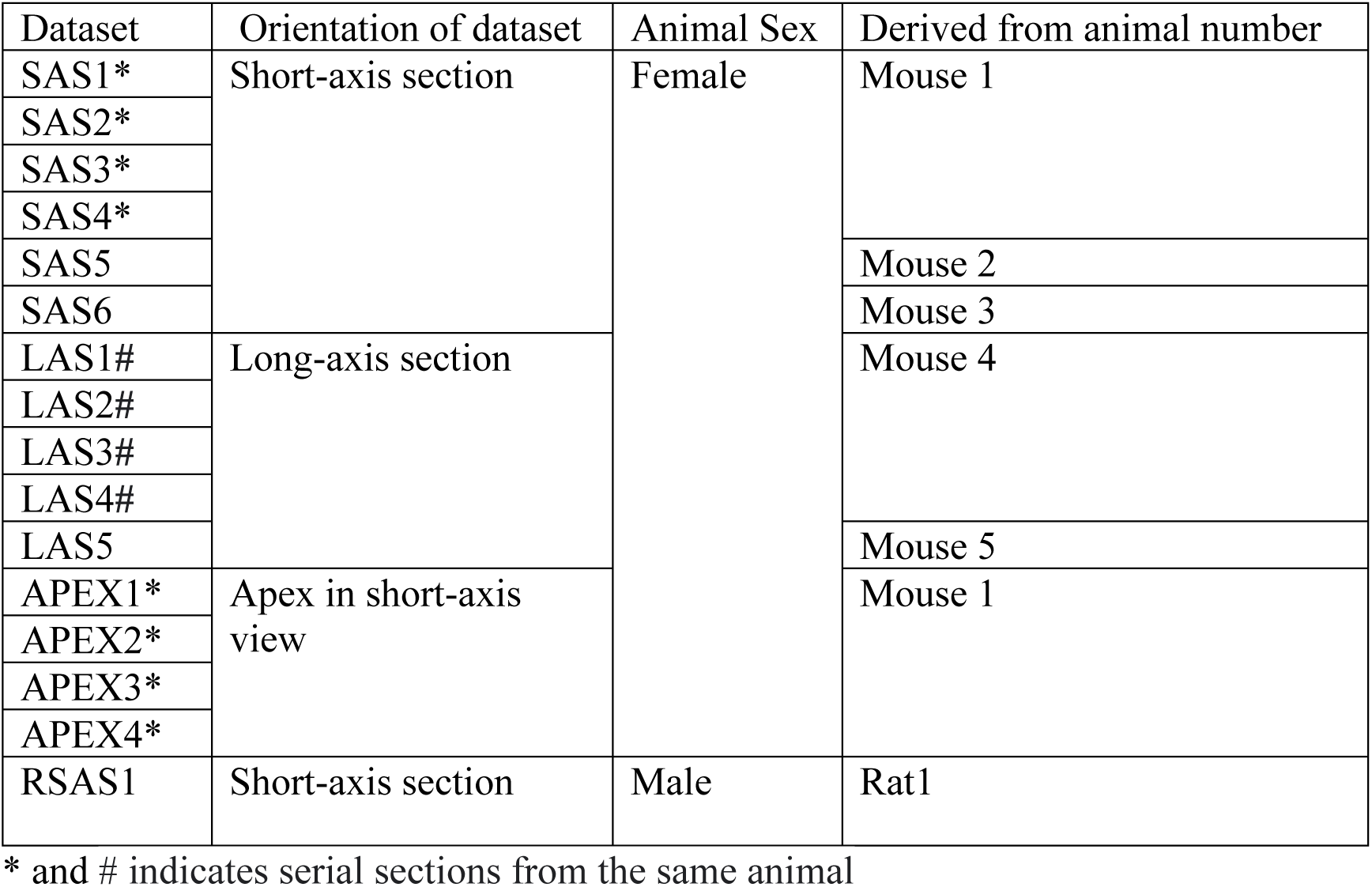
List of sections and animals used in this study.

| Dataset | Orientation of dataset | Animal Sex | Derived from animal number |
| --- | --- | --- | --- |
| SAS1* | Short-axis section | Female | Mouse 1 |
| SAS2* |  |  |  |
| SAS3* |  |  |  |
| SAS4* |  |  |  |
| SAS5 |  |  | Mouse 2 |
| SAS6 |  |  | Mouse 3 |
| LAS1# | Long-axis section |  | Mouse 4 |
| LAS2# |  |  |  |
| LAS3# |  |  |  |
| LAS4# |  |  |  |
| LAS5 |  |  | Mouse 5 |
| APEX1* | Apex in short-axis view |  | Mouse 1 |
| APEX2* |  |  |  |
| APEX3* |  |  |  |
| APEX4* |  |  |  |
| RSAS1 | Short-axis section | Male | Rat1 |

\* and # indicates serial sections from the same animal

## 1 Biological Methods

### 1.1 Experimental procedures

#### CLARITY based clearing protocol applied to the Mouse Heart

All the heart samples used in this study were collected from animals scheduled for culling by the NCBS/inStem Animal Care and Resource Centre facility. We selected a wild-type C57BL/6 strain of female mice aged between 6 weeks and 8 weeks. Once an animal was sacrificed, we immediately performed perfusion to remove blood and clots from the heart tissue. We began by cutting the abdominal cavity of the mouse to access the heart gently. A small incision was made in the right atrium to facilitate fast perfusion of the heart chambers. The perfusion was carried out manually using a 26 Gauge syringe needle inserted at an inclined angle, at the apex region of the right ventricle. We injected 10X phosphate-buffered saline solution (PBS) with a stock solution containing 1.37 M NaCl, 27 mM KCl, 100 mM Na2HPO4, and 18mM KH2PO4, with the pH adjusted to 7.4. Initially, ice cold heparinized 1X PBS was passed through the syringe, followed by ice cold 4% paraformaldehyde (PFA). Subsequently, a hydrogel monomer solution consisting of 4% acrylamide, 4% PFA, 0.5% Bisacrylamide and 0.25% photo-initiator 2, 20-Azobis[2-(2-imidazolin-2-yl)propane] dihydrochloride (VA-044, Wako Chemicals USA) in PBS was perfused through the heart, as described previously for CLARITY based clearing of brain tissue [1]. The fixed mouse heart sample was transferred into a 50ml tube and incubated at 40 degrees celsius for seven days in the hydrogel monomer solution. The fixed heart tissues were then degassed for 10 min using a vacuum chamber at room temperature. To initiate tissue-hydrogel hybridization and polymerization, the processed heart tissues were then incubated for 3 hours at 37 degrees celsius. After polymerization, excess gel material was carefully removed by gently rubbing the tissue with soft tissue wipes. The tissue was transferred to 50 ml tubes for 1X-PBS washes, which were carried out 3 times for a duration of 10 minutes each time. The tissue was further incubated with a clearing buffer (8% SDS and 4% boric acid in 1X PBS (pH 8.5)) for 20-30 days at 37 degrees Celsius, in a shaking incubator (180 rpm) with a buffer exchange occurring every week. This CLARITY based approach applied to the heart tissue samples resulted in transparent tissue (Figure 1A and Supplementary Figure 1A) which could be imaged using a confocal microscope.

### 1.2 Tissue preparation

#### Sectioning

The cleared heart tissue was affixed with superglue at either its short axis or long axis orientation in a specimen tube (Compresstome^®^ VF-300 OZ, Precisionary instruments). The specimen tube was a cylindrical holder with its outer rim fixed and the inside platform (or stage) being movable. The glued heart tissue was embedded in 2.5% low-melting agarose. For a short axis view, the tissue was placed so that the base of the heart was touching the stage of the specimen tube (Supplementary Figure 1B). For a long axis view, the tissue was kept in a plane on the stage, allowing for its four chambers to be seen (Supplementary Figure 1B). We used the compresstome to obtain 500µ*m* thick tissue sections with an oscillation frequency of 7 units and a speed of 1.5 mm/sec. The compresstome blade was kept close to the specimen tube, enabling the compression effect of sectioning to be distributed perpendicular to either the long axis or the short axis of the heart sample. The sectioned tissue was collected in 1X PBS in the chamber associated with the compresstome. Each section was carefully transferred to one of 24 well plates, filled with 1X PBS, while maintaining the sectioning order. For this study, the short axis sections were approximately 3*mm* away from the apex and the long axis sections were approximately 3*mm* from the opposing outer walls of the heart. The apex sections were cut in the short axis plane from the apical tip of the heart.

#### Staining

To carry out staining 500µ*m* processed sections were washed in 1X-PBS thrice over a day and permeabilized using a buffer containing 1% Triton X-100 (H5141, Promega) in PBS (PBST) for one day in a 37*^◦^C* incubator shaker. Subsequently, the tissue sections were incubated in 150µ*g/ml* of Alexa Fluor^TM^ 633 conjugated wheat germ agglutinin (WGA, W21404, ThermoFisher) for one day, to stain cell membranes. The samples were washed with 1X PBS (3 times for 10 minutes each) before incubating them in imaging media (RIMS). For the preparation of RIMS, 40g of Histodenz (Sigma D2158) was dissolved in 30 ml of 0.02 M phosphate buffer with 0.01% sodium azide, pH 7.5, resulting in a final concentration of 88% w/v Histodenz. The labelled tissue samples were incubated in RIMS until the tissue became more transparent [2]. All the staining and washing steps were carried out at room temperature, with gentle shaking. The cleared tissue samples were mounted with fresh RIMS solution using spacers (IS002, SUNjin Lab, Taiwan) of 500µ*m* such that the tissue was sandwiched between coverslips of size 60*mm×* 20*mm*.

### 1.3 3D acquisition of images with a confocal microscope

For image acquisition, we used an Olympus FV3000 microscope. Images were first obtained using a lower magnification objective (Olympus PlanApo 1.25X/ air objective) to image a complete area of the heart tissue section. This low-resolution image was used to map high-resolution imaging areas of interest, using the Olympus fluoView^TM^ software. Then, micron scale imaging was carried out with the Olympus UCPLFN 20X Corr M32 85 mm scale air objective (NA=0.73). We used a 640nm laser line for excitation and FV3000 high sensitivity spectral detectors (gallium arsenide phosphide (GaAsP) photomultiplier tube (PMT)) for detection of emission over a range of 650-670 nm. Each field of view covered approximately 320 *×* 320 pixels, with a voxel size of 1.98 *×* 1.98 *×* 1.98µ*m*^3^ and a depth of *∼* 300µ*m*. Here we under-sampled in the X and Y directions to obtain an isotropic voxel resolution, equivalent to sampling interval in the Z direction. Using the fluoview-map function, we ensured that the acquired 3D images were continuous, and had at least 25% overlap with their respective neighbouring fields of view. To minimize the laser attenuation at deeper regions of the tissue sample, the laser power was corrected (i.e., increased) with the help of the Bright Z function, with a manual judgement based on the quality of the intensity obtained at deeper layers. Each field of view was manually corrected for laser power intensity, increasing this by up to 10% with increased depth. The images were acquired following a snake pattern from row to row. Image reconstructions were performed using computer vision algorithms (as described in Section 2) using custom-written Matlab and C++ scripts. After imaging, the heart tissue samples were stored in RIMS at room temperature and protected from exposure to light.

### 1.4 Alignment of different short-axis sections

We aligned the short axis datasets to ensure uniformity across different short axis sections, using the AHA classification and the capillary vessel description for the PSAX-PML level, used in echocardiogram studies. We used the SAS3 dataset (Table 1) as a reference. First, we aligned the posterior and anterior papillary muscles to the positions of the anterior and inferior regions globally. All other datasets were aligned using the outer wall and papillary muscle morphology to the SAS3 dataset, with the help of a Matlab script and the ImageJ package. To obtain consistent between section alignments at a coarser scale some datasets were reflected (horizontally or vertically) when needed, using ImageJ. For example, the SAS2 and SAS4 samples both required a vertical reflection. The Matlab script we wrote takes the SAS3 dataset as a reference and shows it as a transparent layer. This transparent layer can be rotated by an angular value to allow for fine alignment changes. Once the angular value for in plane rotation had been determined, the short axis datasets were rotated in-plane and saved using ImageJ. The SAS3 and SAS4 had similar morphology. The SAS1 dataset required a -10°in plane rotation and the SAS2 dataset required a -7°in plane rotation. The SAS5 dataset was manually rotated by 180 °in-plane. At the end of this process, all the short axis datasets had a consistent alignment according to the AHA classification, including the positioning of major blood vessels (Supplementary Figure 6).

### 1.5 Analysis of short-axis sections from a rat heart

We also analyzed a four-chambered heart from a different species, that of a wild-type wistar strain of a male rat, approximately 8 weeks in age, which had been scheduled for culling in the NCBS/inStem animal facility. Once the rat had been sacrificed we performed a similar procedure as described above for the mouse hearts. Once the perfusion was completed, the heart was excised and stored in 4%PFA at 40 degrees celsius. To enable the tissue to withstand freezing temperatures, the fixed heart was incubated in 30% sucrose for 5hrs before sectioning. The rat heart was cut into two thick blocks perpendicular to the long axis of the heart (i.e., short-axis-views). The resulting mid-ventricular region was suitable for cryo-sectioning. On the sectioning day, the rat heart was inserted in a mould containing tissue freezing medium and allowed to solidify at *−*200*^◦^C*. The frozen sample was attached to a holder for cryostat (slee mev+), where the heart specimen was placed perpendicular to the long axis of the heart and sectioned into 100µ*m* slices from the midventricular region. The sections were carefully transferred to 24 well plates containing 1X PBS, and were then washed (3 times for 10 minutes each) to remove freezing media. Afterwards, the tissue sections were incubated in 150µ*g/ml* of Alexa FluorTM633 conjugated wheat germ agglutinin (WGA, W21404, ThermoFisher) for one day to stain the cell membrane. The samples were washed with 1X PBS (3 times for 10 minutes each) and incubated in RIMS subsequently for another day. We used a positively charged glass slide for mounting the rat heart tissue sections and custom made 100µ*m* spacers (100µ*m* plastic sheets). The stained heart sections were placed in this glass slide set up with RIMS and sealed with a cover glass in preparation for imaging. We used an Olympus FV3000 microscope and a lower magnification objective (Olympus PlanApo 1.25X/ air objective) to image the complete area of the tissue section to determine the best plane of view. This low-resolution image was used to map the high-resolution imaging area of interest using the Olympus fluoView software. The micron scale imaging was done with the Olympus UCPLFN 20X Corr M32 85 mm scale air objective (NA=0.73). Each field of view consisted of 320 *×* 320 voxels per slice, with a voxel size of 1.98 *×* 1.98 *×* 1.98µ*m*^3^, over a depth of 50µ*m*. As with the mouse hearts, we undersampled in the X, Y directions to obtain isotropic pixels at the resolution of the sampling in the Z direction. Since the area to cover was large, consisting of 300 fields of view in total, we used automatic tile acquisition via the fluoView software platform. Using the fluoview-map function, we ensured that the acquired 3D images were continuous with an overlap of 25% with their neighbouring tiles, regulated by a motorized stage of the microscope. The fields of view were obtained row by row, following a snake pattern, and were then stitched using custom-built software.

### 1.6 Ethics statement

The animal preparation and image work was conducted at the NCBS/inStem Animal Care and Resource Centre and was approved by the inStem Institutional Animal Ethics Committee, following the norms specified by the Committee (INS-IAE-2020/12(N)) for control and supervision of experiments on animals (Government of India). We used the C57BL/6 strain of female mice and these were housed in the institute animal house, and were maintained in a 12 h light/dark cycle. The animals used in our studies were 6-8 weeks in age.

## 2 Computer Vision Methods

### 2.1 Deconvolution of acquired data

Since the confocal images were acquired in three dimensions the resulting data was blurred in a manner that depended on the shape of the point spread function (PSF) of the microscope. A pseudo-color image of PSF generated based on our microscope settings is shown in Supplementary Information Figure (SI-Fig.) 1. To mitigate the effects of this blur we deconvolved each tile using an iterative Richardson-Lucy (RL) deconvolution method, with Total variation (TV) regularization (RL-TV) [3]. The algorithm minimized the following objective function:

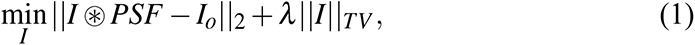

where ⊛ represents the convolution operation and TV is the total variation norm. In our implementation we set λ = 0.01 and processed each field of view for 20 iterations.

### 2.2 Denoising

As depth in the tissue increased, the signal to noise ratio decreased. As a result, the visual quality of the deeper layers was poorer than that of the shallow layers. To mitigate this effect we applied an unsupervised dictionary based method for denoising the images following the deconvolution stage. The method was based on the assumption that layers in the tissue are self similar so that the ultrastructure of the tissue is similar in different depth layers of a single field of view. We trained a sparse (*m* = 256) element 2D dictionary of patches of size 16 *×* 16 using the sparse dictionary learning method of [4]. The shallow layers were relatively free from both depth and other optical degradation effects. The dictionary (*D*) was learned from data samples (*x_i_*) from the shallow layers using the alternating minimization approach in [5]. The method involved alternating between between fixing *D* and solving the resulting basis pursuit denoising problem in Eq. (2) below

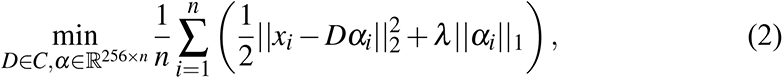

followed by fixing α and updating the dictionary *D* using coordinate descent. Here, 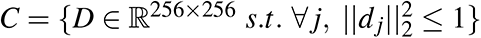, and α*_i_* are the sparse codes corresponding to data element *x_i_*. We used a value of λ = 0.15 as the regularization parameter in our experiments and ran the optimization for 1000 iterations. The set of learned dictionary patches using tissue samples from the SAS1 dataset is shown in SI-Fig. 2. The images acquired from deeper layers in the tissue sample can then be denoised using this learned sparse dictionary. To accomplish this, at each voxel in a degraded image we constructed a 16 *×* 16 patch centered at the voxel and estimated a denoised patch α by solving the sparse coding problem in Eq. (3) using a lars/homotopy method [6]:

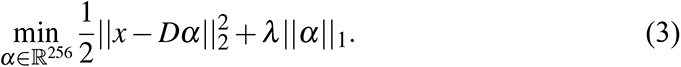

The final denoised image was reconstructed as an average of the denoised patches of the overlapping windows at each voxel. We learned a separate dictionary for each field of view so that the structure in one field of view did not affect the reconstruction in another.

**SI-Fig. 1:**
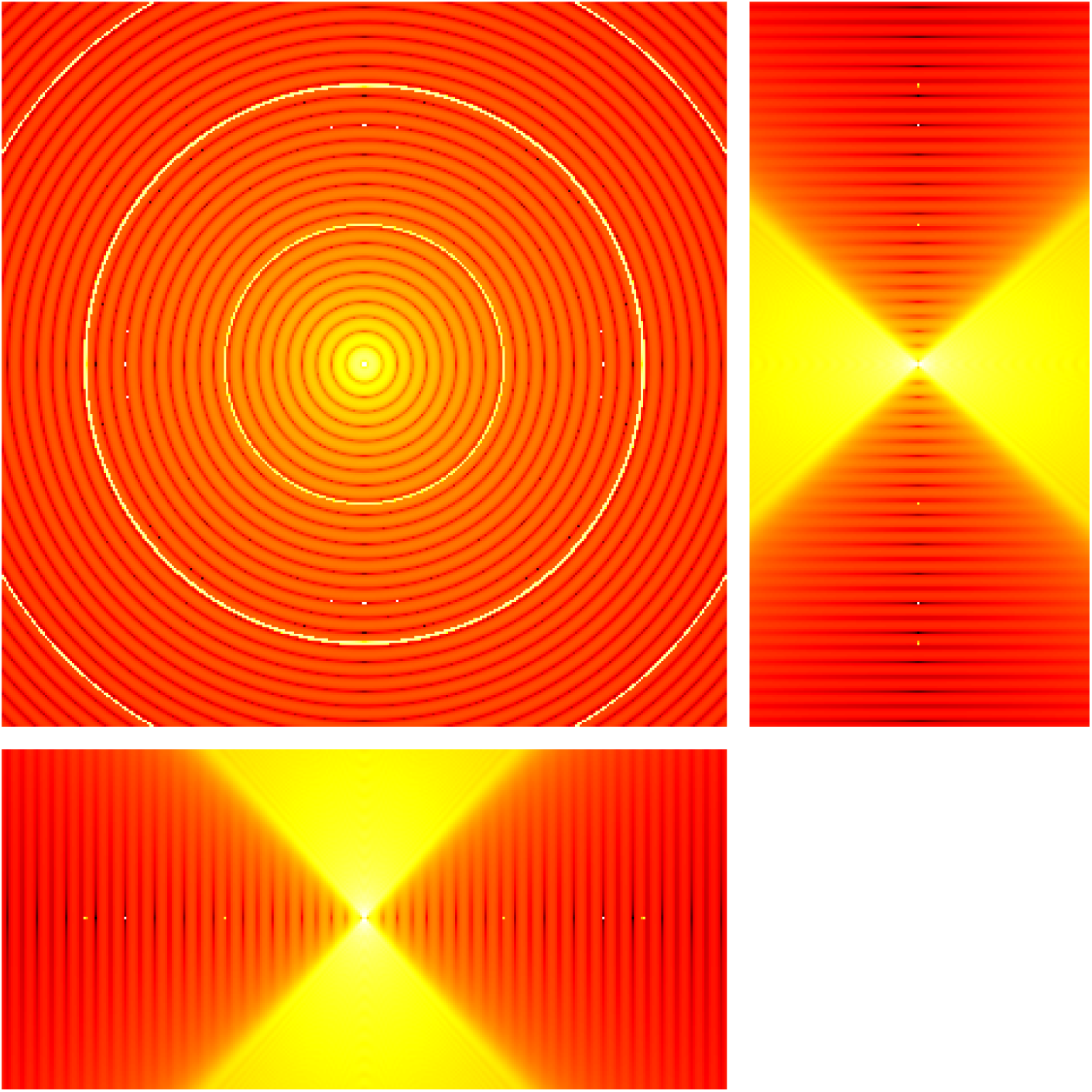
A visualization of the middle planes of a 3D point spread function (PSF) along the *XY* (top left), *XZ* (bottom left) and *YZ* (top right) directions, shown in log intensity scale (increasing from black through yellow and red to white).

**SI-Fig. 2:**
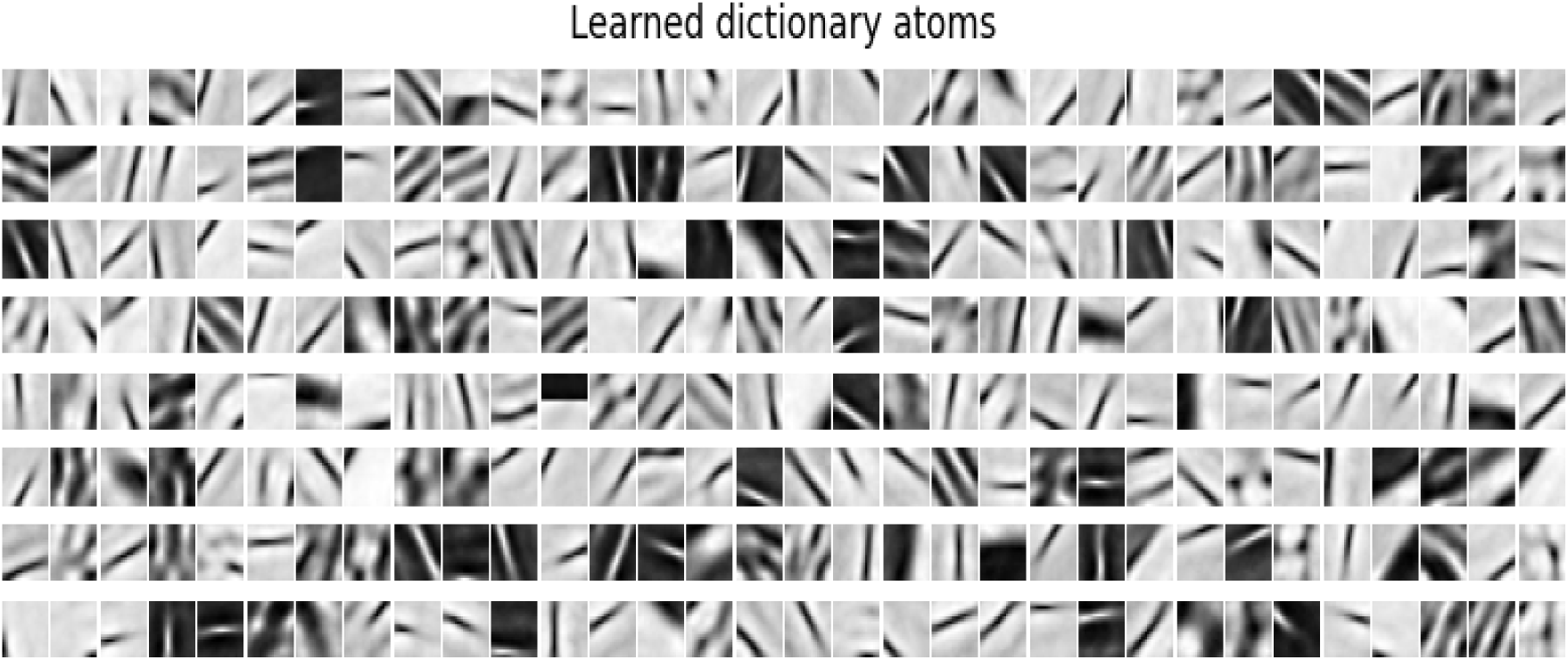
A visualization of the dictionary atoms learned for the mouse cardiac tissue microscopy image dataset SAS1.

### 2.3 Stitching 3D blocks

Each tissue section was too large to be imaged at once so we imaged multiple square shaped fields of view (tiles) of 320 *×* 320 isotropic voxels of length 1.98µ*m* in each dimension, in the regions containing tissue samples. Adjacent tiles were set to have an overlap of 12.5% in every direction. We used the image registration method described in [7] for regions with valid data. The method involved a two stage registration process, with a local pairwise registration followed by a global registration. In the first local registration stage, we started with an initial guess for the location of each tile, derived from the microscope stage settings and assumed a 40 voxel (12.5%) overlap value. For every pair of adjacent tiles (a, b) we used the maximum phase correlation[8] based registration to estimate the relative shift, *p_ab_* between the pair.

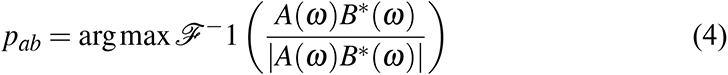

where, *ℱ^−^*^1^ represents the inverse Fourier transform and *A*(ω) and *B^∗^*(ω) are the Fourier transform and the complex conjugate of the Fourier transform of tile a and tile b, respectively. For every imaged tile, a shift was computed with each of its 4-neighbours in the 2D imaging plane. This local pairwise registration process resulted in a refined list of (*p_ab_*) pairwise relative shift values.

In the second global registration stage, a global optimal tile location of each tile (*p_a_, p_b_, …*) was computed with respect to the top left corner of the image. In all our datasets this corner tile was empty and was only used to define a common reference frame. The vector of optimal tile positions *P* for all tiles *T* = *{a, b, ..}* was then given by

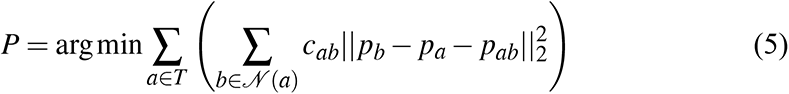

where *c_ab_* was the correlation value between the pair *a, b*. This global registration was accomplished by solving an over-determined system of sparse linear equations for position [9]. This was done by iteratively eliminating all pairs of outlier pairwise distances. A local shift was labelled an outlier if it was over three standard deviations away from the mean shift.

SI-Figures 2B and 2C illustrate the stitching process for a short axis section of a mouse heart. Two sample tiles are demarcated by red and blue bounding boxes.

### 2.4 Orientation field estimation

The orientation field was estimated at each voxel in the stitched and denoised image stack. We used the structure tensor[10] *s* = *G*_ρ_ ⊛(∇_σ_ *I*)(∇_σ_ *I*)*^T^*, where *G*_ρ_ is a Gaussian with standard deviation ρ, ⊛ is the convolution operation and ∇_σ_ represents the intensity gradient at a Gaussian smoothing scale of σ. We used a noise scale σ = 0.5 and feature scale of ρ = 3 voxel units. The orientation was then set to align with the eigenvector of the structure tensor corresponding to the eigenvalue with smallest magnitude.

### 2.5 Computation of the Helix Angle

The helix angle α*H* is typically defined in a manner that is relative to the local direction normal to the outer heart wall. To estimate the wall normal we computed a single pixel wide boundary of the heart in a short axis section and fit a circle tangential at each point along the outer boundary, using 80 sample points along the boundary in each direction. The direction of the heart wall normal was then associated with the inward radial vector of the circular fit (SI-Figure 4-A). The local helix angle α*_H_*, as illustrated in Figure 2-A, was calculated using the projection of the orientation onto the tangential plane defined by the heart wall normal. The angle varied from *−*90*^◦^* to 90*^◦^*, with 0*^◦^* representing the in plane circumferential fibers and *±*90*^◦^* representing fibers pointing out of the short axis plane, in the long axis direction of the heart.

### 2.6 Smoothing the Estimated Orientation Field

Given the thickness of tissue samples used in our study, imaging data in deeper layers, where light penetration was reduced, was noisy. In addition, optical factors including light scattering, photo bleaching and optical aberrations in the microscope lens, also diminished the image quality. To mitigate the affect of the reduced image quality on orientation estimation we averaged the orientation field over small neighborhoods. Whereas orientations are directionless, their representation using the eigenvector with the smallest eigenvalue of the structure tensor is not. Two vectors whose components have the same magnitude but differ in signs represent the same orientation, so these direction vectors cannot be directly averaged component wise.

To smooth the orientations we first computed the rank-1 tensor constructed as *s* = *uu^T^*, where *s* is a 3 *×* 3 matrix and *u* was the local unit direction vector. This rank-1 tensor was invariant to flips, since *s* = *uu^T^* = (*−u*)(*−u*)*^T^* . In fact, *s ∈ Gr*(3, 1), represents the Grassmann manifold of one dimensional subspaces (lines) in 3 dimensional Euclidean space. While it was possible to use the weighted Karcher mean to smooth the resulting tensors component-wise, an iterative approach to doing so was slow and did not scale well for to handle large volumes of data. We therefore opted for an approximate strategy. We averaged the orientation tensors *s* component-wise using a local weighted average, and then projected back to the space of direction vectors. Specifically, the smoothed tensor *s*^ at any location **x** was given by

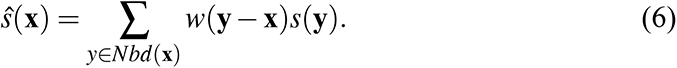

Here, *w*(*·*) is a scalar weight, which was empirically chosen to be a Gaussian with σ = 4 as defined below. The smoothed direction vector *u*^ at **x** was then given by the eigenvector of the *s*^ matrix corresponding to the eigenvalue with the largest magnitude . We only carried out smoothing in regions within the heart tissue by setting the weight *w*(**z**) to zero in regions with missing data:

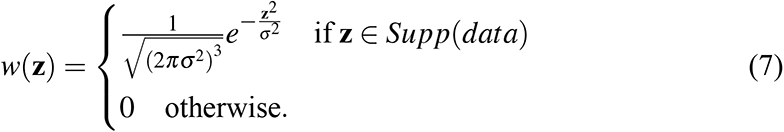

### 2.7 Φ, θ and α*H* calculation and colormap generation

The colormaps for Φ, θ and α*H* were generated using the smoothed orientation field. We calculated the Φ and θ angles at each voxel with valid data and then mapped these angles to color values using a linear scale in the parula colormap.

To obtain the α*H* colormaps we considered voxels with valid data and then computed the average value of the helix angle over all radial penetration lines overlapping at the voxel. A penetration line at a point (*x, y*) in a short axis plane was considered to overlap an integer valued voxel (*i, j*) when *i ≤ x ≤ i* + 1 and *j ≤ y ≤ j* + 1. Due to irregularities in boundary shape, which in turn affected the estimate of the heart wall normal, it was possible for a few isolated voxels, containing valid orientation estimates, to have no penetration line passing through them. The α*H* value at each such location was set to the average α*H* value over a 3 *×* 3 voxel neighbourhood.

### 2.8 3D Rendered Visualizations and Animations

Orientation glyphs and streamlines were procedurally generated and rendered using a custom Python module within the open-source animation software Blender. For each sample volume, an orientation field and a tiff stack representing the WGA tissue staining were imported as N-dimensional NumPy arrays.

#### Glyphs

The orientation field was represented by a 3-dimensional array of rotated cylinders, referred to as orientation glyphs. The input vector field was approximated by a 3-dimensional grid of equally spaced vertices, downsampled such that the number of vertices was not larger than 75,000. At each vertex of the downsampled grid, a cylinder primitive shape was created, and rotated proportionally to the components of the vector field at the vertex position. A parula colormap was applied to the cylinders proportional to the Phi angle, defined as the arc cosine of the absolute value of the z component of the vector field.

#### Streamlines

Bidirectional streamlines were represented as curves extruded from polylines, whose points were computed as follows. A set of up to 25,000 points were selected from a random sample of voxels from the vector field, and each sample voxel location was the initial point for a streamline. Since the direction of the vector field represents the orientation of the tissue, and the sign of the orientation does not matter, each starting point initialized both a positive and negative streamline. Both positive and negative streamlines originated from a single starting point, and were grown by iteratively adding new points along the polyline. For each iteration, the location of the next point was calculated by adding a displacement proportional to the vector field and the current point, multiplied by the sign, such that the positive streamline is displaced by the positive value of the field, and the negative streamline by the negative value of the field. At each new position, the value of the field at that voxel acts to determine the position of the following point along the polyline. A parula colormap was applied to the streamlines in proportion to the Phi angle at the streamlines starting point within the vector field. To avoid artifacts arising where the streamlines extend beyond tissue, a binary mask of the tissue volume with the same dimensions as the vector field served as a boundary condition.

## References

1. Peter Libby, MD, PhD and Douglas P. Zipes, MD; Edited by Robert O. Bonow, MD, MS, Douglas L. Mann, MD, Gordon F. Tomaselli, MD and Deepak Bhatt, Braunwald’s Heart Disease: A Textbook of Cardiovascular Medicine, 2-Volume Set (ed. 11, 2018; https://www.uk.elsevierhealth.com/braunwalds-heart-disease-a-textbook-of-cardiovascular-medicine-2-volume-set-9780323463423.html).

2. S. H. Gilbert, A. P. Benson, P. Li, A. V. Holden, Regional localisation of left ventricular sheet structure: integration with current models of cardiac fibre, sheet and band structure. Eur. J. Cardio-Thorac. Surg. Off. J. Eur. Assoc. Cardio-Thorac. Surg. 32, 231–249 (2007).

3. T. Aumentado-Armstrong, A. Kadivar, P. Savadjiev, S. W. Zucker, K. Siddiqi, Conduction in the Heart Wall: Helicoidal Fibers Minimize Diffusion Bias. Sci. Rep. 8, 7165 (2018).

4. R. J. Young, A. V. Panfilov, Anisotropy of wave propagation in the heart can be modeled by a Riemannian electrophysiological metric. Proc. Natl. Acad. Sci. U. S. A. 107, 15063–15068 (2010).

5. C. von Deuster, E. Sammut, L. Asner, D. Nordsletten, P. Lamata, C. T. Stoeck, S. Kozerke, R. Razavi, Studying Dynamic Myofiber Aggregate Reorientation in Dilated Cardiomyopathy Using In Vivo Magnetic Resonance Diffusion Tensor Imaging. Circ. Cardiovasc. Imaging. 9, e005018 (2016).

6. J. Chen, S.-K. Song, W. Liu, M. McLean, J. S. Allen, J. Tan, S. A. Wickline, X. Yu, Remodeling of cardiac fiber structure after infarction in rats quantified with diffusion tensor MRI. Am. J. Physiol.-Heart Circ. Physiol. 285, H946–H954 (2003).

7. L. Geerts-Ossevoort, P. Bovendeerd, F. Prinzen, T. Arts, K. Nicolay, in Computers in Cardiology 2001*. Vol.*28 (Cat. No.01CH37287) (2001), pp. 621–624.

8. R. H. Anderson, M. Smerup, D. Sanchez-Quintana, M. Loukas, P. P. Lunkenheimer, The three-dimensional arrangement of the myocytes in the ventricular walls. Clin. Anat. N. Y. N. 22, 64–76 (2009).

9. P. Helm, M. F. Beg, M. I. Miller, R. L. Winslow, Measuring and mapping cardiac fiber and laminar architecture using diffusion tensor MR imaging. Ann. N. Y. Acad. Sci. 1047, 296–307 (2005).

10. C. S. Peskin, Fiber architecture of the left ventricular wall: An asymptotic analysis. Commun. Pure Appl. Math. 42, 79–113 (1989).

11. A. Horowitz, M. Perl, S. Sideman, Geodesics as a mechanically optimal fiber geometry for the left ventricle. Basic Res. Cardiol. 88 **Suppl 2**, 67–74 (1993).

12. P. Savadjiev, G. J. Strijkers, A. J. Bakermans, E. Piuze, S. W. Zucker, K. Siddiqi, Heart wall myofibers are arranged in minimal surfaces to optimize organ function. Proc. Natl. Acad. Sci. 109, 9248–9253 (2012).

13. T. Seidel, J.-C. Edelmann, F. B. Sachse, Analyzing Remodeling of Cardiac Tissue: A Comprehensive Approach Based on Confocal Microscopy and 3D Reconstructions. Ann. Biomed. Eng. 44, 1436–1448 (2016).

14. I. Nehrhoff, J. Ripoll, R. Samaniego, M. Desco, M. V. Gómez-Gaviro, Looking inside the heart: a see-through view of the vascular tree. Biomed. Opt. Express. 8, 3110– 3118 (2017).

15. F. Perbellini, A. K. L. Liu, S. A. Watson, I. Bardi, S. M. Rothery, C. M. Terracciano, Free-of-Acrylamide SDS-based Tissue Clearing (FASTClear) for three dimensional visualization of myocardial tissue. Sci. Rep. 7, 5188 (2017).

16. A. A. Young, I. J. Legrice, M. A. Young, B. H. Smaill, Extended confocal microscopy of myocardial laminae and collagen network. J. Microsc. 192, 139–150 (1998).

17. G. H. Granlund, H. Knutsson, Signal Processing for Computer Vision (Springer Science & Business Media, 1994).

18. J. Chen, W. Liu, H. Zhang, L. Lacy, X. Yang, S.-K. Song, S. A. Wickline, X. Yu, Regional ventricular wall thickening reflects changes in cardiac fiber and sheet structure during contraction: quantification with diffusion tensor MRI. Am. J. Physiol.-Heart Circ. Physiol. 289, H1898–H1907 (2005).

19. H. M. Spotnitz, Macro design, structure, and mechanics of the left ventricle. J. Thorac. Cardiovasc. Surg. 119, 1053–1077 (2000).

20. F. Poveda, D. Gil, E. Martí, A. Andaluz, M. Ballester, F. Carreras, Helical Structure of the Cardiac Ventricular Anatomy Assessed by Diffusion Tensor Magnetic Resonance Imaging With Multiresolution Tractography. Rev. Esp. Cardiol. Engl. Ed. 66, 782–790 (2013).

21. M. F. Beg, P. A. Helm, E. McVeigh, M. I. Miller, R. L. Winslow, Computational cardiac anatomy using MRI. Magn. Reson. Med. 52, 1167–1174 (2004).

22. J.-M. Peyrat, M. Sermesant, X. Pennec, H. Delingette, C. Xu, E. R. McVeigh, N. Ayache, A computational framework for the statistical analysis of cardiac diffusion tensors: application to a small database of canine hearts. IEEE Trans. Med. Imaging. 26, 1500–1514 (2007).

23. H. Lombaert, J.-M. Peyrat, P. Croisille, S. Rapacchi, L. Fanton, F. Cheriet, P. Clarysse, I. Magnin, H. Delingette, N. Ayache, Human atlas of the cardiac fiber architecture: study on a healthy population. IEEE Trans. Med. Imaging. 31, 1436– 1447 (2012).

24. D. Sedmera, R. G. Gourdie, Why do we have Purkinje fibers deep in our heart? Physiol. Res. 63, S9–18 (2014).

25. D. Romero, O. Camara, F. Sachse, R. Sebastian, Analysis of Microstructure of the Cardiac Conduction System Based on Three-Dimensional Confocal Microscopy. PLOS ONE. 11, e0164093 (2016).

26. N. E. Diamant, Physiology of Esophageal Motor Function. Gastroenterol. Clin. North Am. 18, 179–194 (1989).

27. R. K. Mittal, Longitudinal muscle of the esophagus: its role in esophageal health and disease. Curr. Opin. Gastroenterol. 29, 421–430 (2013).

28. M. L. Scimone, L. E. Cote, P. W. Reddien, Orthogonal muscle fibres have different instructive roles in planarian regeneration. Nature. 551, 623–628 (2017).

29. R. Tomer, L. Ye, B. Hsueh, K. Deisseroth, Advanced CLARITY for rapid and high-resolution imaging of intact tissues. Nat. Protoc. 9, 1682–1697 (2014).

30. N. Dey, L. Blanc-Feraud, C. Zimmer, P. Roux, Z. Kam, J.-C. Olivo-Marin, J. Zerubia, Richardson–Lucy algorithm with total variation regularization for 3D confocal microscope deconvolution. Microsc. Res. Tech. 69, 260–266 (2006).

31. J. Mairal, F. Bach, J. Ponce, Sparse Modeling for Image and Vision Processing (2014) (available at https://arxiv.org/abs/1411.3230v2).

32. D. Zukić, M. Jackson, D. Dimiduk, S. Donegan, M. Groeber, M. McCormick, ITKMontage: A Software Module for Image Stitching. Integrating Mater. Manuf. Innov. 10, 115–124 (2021).

33. H. Knutsson, C.-F. Westin, M. Andersson, in Image Analysis, A. Heyden, F. Kahl, Eds. (Springer, Berlin, Heidelberg, 2011), pp. 545–556.

34. M. D. Cerqueira, N. J. Weissman, V. Dilsizian, A. K. Jacobs, S. Kaul, W. K. Laskey, D. J. Pennell, J. A. Rumberger, T. Ryan, M. S. Verani, American Heart Association Writing Group on Myocardial Segmentation and Registration for Cardiac Imaging, Standardized myocardial segmentation and nomenclature for tomographic imaging of the heart. A statement for healthcare professionals from the Cardiac Imaging Committee of the Council on Clinical Cardiology of the American Heart Association. Circulation. 105, 539–542 (2002).

## References

[1] Kwanghun Chung, et al. “Structural and molecular interrogation of intact biological systems”. In: Nature 497.7449 (2013), pp. 332–337.

[2] Bin Yang, et al. “Single-cell phenotyping within transparent intact tissue through whole-body clearing”. In: Cell 158.4 (2014), pp. 945–958.

[3] Nicolas Dey, et al. “Richardson–Lucy algorithm with total variation regularization for 3D confocal microscope deconvolution”. In: Microscopy research and technique 69.4 (2006), pp. 260–266.

[4] Julien Mairal, Francis Bach, Jean Ponce, et al. “Sparse modeling for image and vision processing”. In: Foundations and Trends® in Computer Graphics and Vision 8.2-3 (2014), pp. 85–283.

[5] Julien Mairal, et al. “Online dictionary learning for sparse coding”. In: Proceedings of the 26th annual international conference on machine learning. 2009, pp. 689–696.

[6] Iddo Drori and David L Donoho. “Solution of L1 minimization problems by LARS/homotopy methods”. In: 2006 IEEE International Conference on Acoustics Speech and Signal Processing Proceedings. Vol. 3. IEEE. 2006, pp. III–III.

[7] Stephan Preibisch, Stephan Saalfeld, and Pavel Tomancak. “Globally optimal stitching of tiled 3D microscopic image acquisitions”. In: Bioinformatics 25.11 (2009), pp. 1463–1465.

[8] C.D. Kuglin and D.C. Hines. “The phase correlation image alignment method”. In: Proc. Int. Conf. on Cybernetics and Society. IEEE, Sept. 1975, pp. 163–165.

[9] Dženan Zukić, et al. “ITKMontage: A Software Module for Image Stitching”. In: Integrating Materials and Manufacturing Innovation 10.1 (2021), pp. 115–124.

[10] Hans Knutsson, Carl-Fredrik Westin, and Mats Andersson. “Representing local structure using tensors II”. In: Scandinavian conference on image analysis. Springer. 2011, pp. 545–556.

